# The Invariance Hypothesis Implies Domain-Specific Regions in Visual Cortex

**DOI:** 10.1101/004473

**Authors:** Joel Z Leibo, Qianli Liao, Fabio Anselmi, Tomaso Poggio

## Abstract

Is visual cortex made up of general-purpose information processing machinery, or does it consist of a collection of specialized modules? If prior knowledge, acquired from learning a set of objects is only transferable to new objects that share properties with the old, then the recognition system’s optimal organization must be one containing specialized modules for different object classes. Our analysis starts from a premise we call the invariance hypothesis: that the computational goal of the ventral stream is to compute an invariant-to-transformations and discriminative signature for recognition. The key condition enabling approximate transfer of invariance without sacrificing discriminability turns out to be that the learned and novel objects transform similarly. This implies that the optimal recognition system must contain subsystems trained only with data from similarly-transforming objects and suggests a novel interpretation of domain-specific regions like the fusiform face area (FFA). Furthermore, we can define an index of transformation-compatibility, computable from videos, that can be combined with information about the statistics of natural vision to yield predictions for which object categories ought to have domain-specific regions in agreement with the available data. The result is a unifying account linking the large literature on view-based recognition with the wealth of experimental evidence concerning domain-specific regions.

## 1 Introduction

The discovery of category-selective patches in the ventral stream—e.g., the fusiform face area (FFA)—is one of the most robust experimental findings in visual neuroscience [1, 2, 3, 4, 5, 6]. It has also generated significant controversy. From a computational perspective, much of the debate hinges on the question of whether the algorithm implemented by the ventral stream requires subsystems or modules dedicated to the processing of a single class of stimuli [7, 8]. The alternative account holds that visual representations are distributed over many regions [9, 10], and the clustering of category selectivity is not, in itself, functional. Instead, it arises from the interaction of biological constraints like anatomically fixed inter-region connectivity and competitive plasticity mechanisms [11, 12] or the center-periphery organization of visual cortex [13, 14, 15, 16, 17].

The interaction of three factors is thought to give rise to properties of the ventral visual pathway: (1) The computational task; (2) constraints of anatomy and physiology; and (3) the statistics of the visual environment [18, 19, 20, 21, 22]. Differing presuppositions concerning their relative weighting lead to quite different models of the origin of category-selective regions. If the main driver is thought to be the visual environment (factor 3), then perceptual expertise-based accounts of category selective regions are attractive [23, 24, 25]. Alternatively, mechanistic models show how constraints of the neural “hardware” (factor 2) could explain category selectivity [26, 12, 27]. Contrasting with both of these, the perspective of the present paper is one in which computational factors are the main reason for the clustering of category-selective neurons.

The lion’s share of computational modeling in this area has been based on factors 2 and 3. These models seek to explain category selective regions as the inevitable outcome of the interaction between functional processes; typically competitive plasticity, wiring constraints, e.g., local connectivity, and assumptions about the system’s inputs [26, 28, 12, 27]. Mechanistic models of category selectivity may even be able to account for the neuropsychology [29, 30] and behavioral [31, 32] results long believed to support modularity.

**Figure 1:**
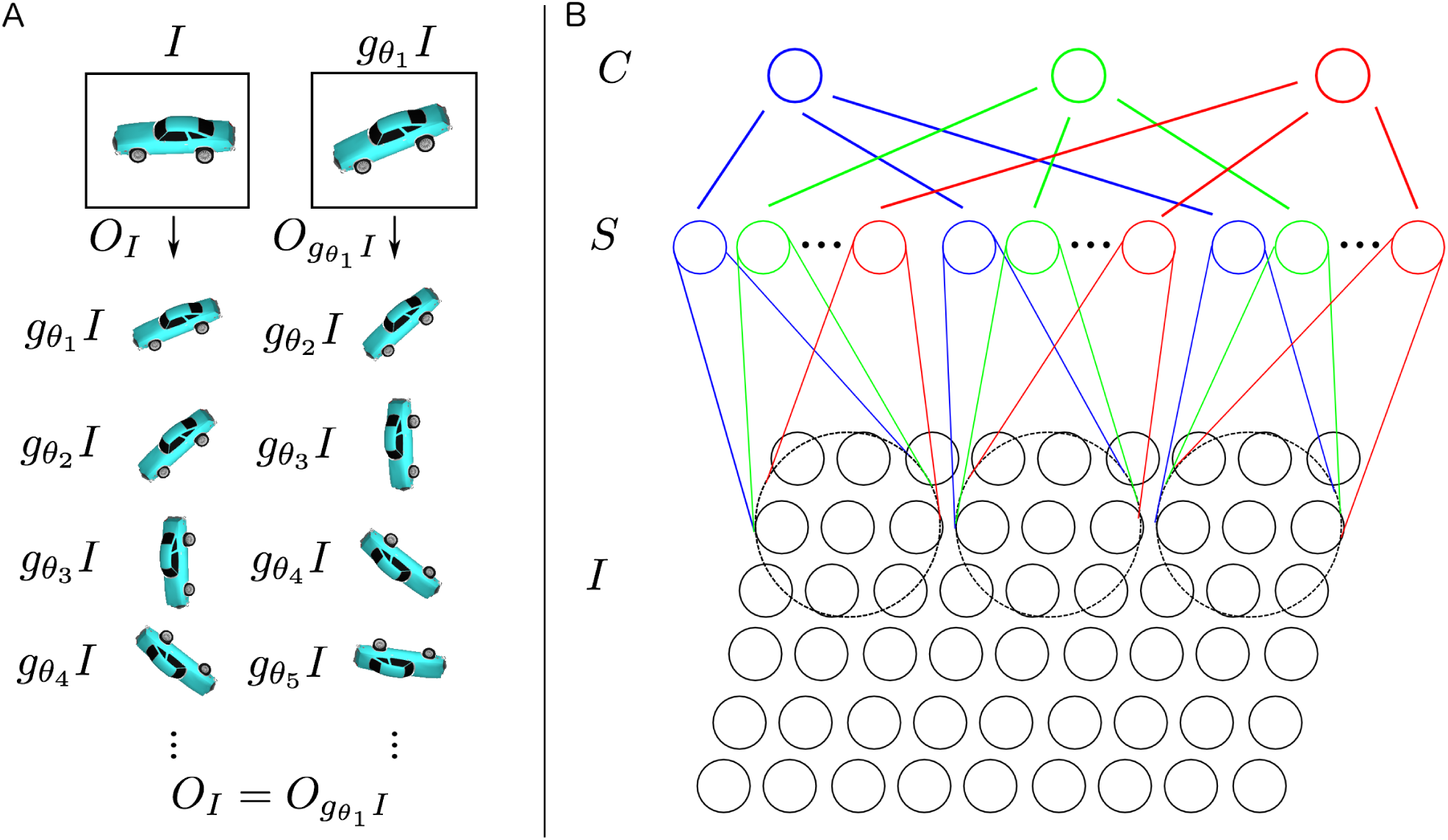
A. The orbit with respect to in-plane rotation is invariant and unique. **B.** Three HW-modules are shown. In this example, each HW-module pools over a 9 *×* 3 region of the image. Each S-unit stores a 9 *×* 3 template and there are three S-units per HW-module.

Another line of evidence seems to explain away the category selective regions. The large-scale topography of object representation is reproducible across subjects [33]. For instance, the scene-selective parahippocampal place area (PPA) is consistently medial to the FFA. To explain this remarkable reproducibility, it has been proposed that the center-periphery organization of early visual areas extends to the later object-selective regions of the ventral stream [13, 14, 15, 17]. In particular, the FFA and other face-selective region are associated with an extension of the central representation, and PPA with the peripheral representation. Consistent with these findings, it has also been argued that real-world size is the organizing principle [16]. Larger objects, e.g., furniture, evoke more medial activation while smaller objects, e.g., a coffee mug, elicit more lateral activity.

Could category selective regions be explained as a consequence of the topography of visual cortex? Both the eccentricity [15] and real-world size [16] hypotheses correctly predict that houses and faces will be represented at opposite ends of the medial-lateral organizing axis. Since eccentricity of presentation is linked with acuity demands, the differing eccentricity profiles across object categories may be able to explain the clustering. However, such accounts offer no way of interpreting macaque results indicating multi-stage processing hierarchies [34, 17]. If clustering was a secondary effect driven by acuity demands, then it would be difficult to explain why, for instance, the macaque face-processing system consists of a hierarchy of patches that are preferentially connected with one another [35].

In macaques, there are 6 discrete face-selective regions in the ventral visual pathway, one posterior lateral face patch (PL), two middle face patches (lateral-ML and fundus-MF), and three anterior face patches, the anterior fundus (AF), anterior lateral (AL), and anterior medial (AM) patches [2, 36]. At least some of these patches are organized into a feedforward hierarchy. Visual stimulation evokes a change in the local field potential *∼* 20 ms earlier in ML/MF than in patch AM [34]. Consistent with a hierarchical organization involving information passing from ML/MF to AM via AL, electrical stimulation of ML elicits a response in AL and stimulation in AL elicits a response in AM [35]. In addition, spatial position invariance increases from ML/MF to AL, and increases further to AM [34] as expected for a feedforward processing hierarchy. The firing rates of neurons in ML/MF are most strongly modulated by face viewpoint. Further along the hierarchy, in patch AM, cells are highly selective for individual faces and collectively provide a representation of face identity that tolerates substantial changes in viewpoint [34].

Freiwald and Tsao argued that the network of face patches is *functional*. Response patterns of face patch neurons are consequences of the role they play in the algorithm implemented by the ventral stream. Their results suggest that the face network computes a representation of faces that is—as much as possible—invariant to 3D rotation-in-depth (viewpoint), and that this representation may underlie face identification behavior [34].

We carry out our investigation within the framework provided by a recent theory of invariant object recognition in hierarchical feedforward architectures [37]. It is broadly in accord with other recent perspectives on the ventral stream and the problem of object recognition [38, 22]. The full theory has implications for many outstanding questions that are not directly related to the question of domain specificity we consider here. In other work, it has been shown to yield predictions concerning the cortical magnification factor and visual crowding [39]. It has also been used to motivate novel algorithms in computer vision and speech recognition that perform competitively with the state-of-the-art on difficult benchmark tasks [40, 41, 42, 43, 44]. The same theory, with the additional assumption of a particular Hebbian learning rule, can be used to derive qualitative receptive field properties. The predictions include Gabor-like tuning in early stages of the visual hierarchy [45, 46] and mirror-symmetric orientation tuning curves in the penultimate stage of a face-specific hierarchy computing a view-tolerant representation (as in [34]) [46]. A full account of the new theory is outside the scope of the present work; we refer the interested reader to the references—especially [37] for details.

Note that the theory only applies to the first feedforward pass of information, from the onset of the image to the arrival of its representation in IT cortex approximately 100 ms later. For a recent review of evidence that the feedforward pass computes invariant representations, see [22]. For an alternative perspective, see [11]. Though note also, contrary to a claim in that review, position dependence is fully compatible with the class of models we consider here (including HMAX). [47, 39] explicitly model eccentricity dependence in this framework.

Our account of domain specificity is motivated by the following questions: How can past visual experience be leveraged to improve future recognition of novel individuals? Is any past experience useful for improving at-a-glance recognition of any new object? Or perhaps past experience only transfers to similar objects? Could it even be possible that past experience with certain objects actually impedes the recognition of others?

The invariance hypothesis holds that the computational goal of the ventral stream is to compute a representation that is unique to each object and invariant to identity-preserving transformations. If we accept this premise, the key question becomes: Can transformations learned on one set of objects be reliably transferred to another set of objects? For many visual tasks, the variability due to transformations in a single individual’s appearance is considerably larger than the variability between individuals. These tasks have been called “subordinate level identification” tasks, to distinguish them from between-category (basic-level) tasks. Without prior knowledge of transformations, the subordinate-level task of recognizing a novel individual from a single example image is hopelessly under-constrained.

The main thrust of our argument—to be developed below—is this: The ventral stream computes object representations that are invariant to transformations. Some transformations are *generic*; the ventral stream could learn to discount these from experience with any objects. Translation and scaling are both generic (all 2D affine transformations are). However, it is also necessary to discount many transformations that do not have this property. Many common transformations are not generic; 3D-rotation-in-depth is the primary example we consider here (see Text S1 for more examples). It is not possible to achieve a perfectly view-invariant representation from one 2D example. Out-of-plane rotation depends on information that is not available in a single image, e.g. the object’s 3D structure. Despite this, approximate invariance can still be achieved using prior knowledge of how similar objects transform. In this way, approximate invariance learned on some members of a visual category can facilitate the identification of unfamiliar category members. But, this transferability only goes so far.

Under this account, the key factor determining which objects could be productively grouped together in a domain-specific subsystem is their transformation compatibility. We propose an operational definition that can be computed from videos of transforming objects. Then we use it to explore the question of why certain object classes get dedicated brain regions, e.g., faces and bodies, while others (apparently) do not.

We used 3D graphics to generate a library of videos of objects from various categories undergoing rotations in depth. The model of visual development (or evolution) we consider is highly stylized and non-mechanistic. It is just a clustering algorithm based on our operational definition of transformation compatibility. Despite its simplicity, using the library of depth-rotation videos as inputs, the model predicts large clusters consisting entirely of faces and bodies.

The other objects we tested—vehicles, chairs, and animals—ended up in a large number of small clusters, each consisting of just a few objects. This suggests a novel interpretation of the lateral occipital complex (LOC). Rather than being a “generalist” subsystem, responsible for recognizing objects from diverse categories, our results are consistent with LOC actually being a heterogeneous region that consists of a large number of domain-specific regions too small to be detected with fMRI.

These considerations lead to a view of the ventral visual pathway in which category-selective regions implement a modularity of *content* rather than *process* [48, 49]. Our argument is consistent with process-based accounts, but does not require us to claim that faces are automatically processed in ways that are inapplicable to objects (e.g., gaze detection or gender detection) as claimed by [11]. Nor does it commit us to claiming there is a region that is specialized for the process of subordinate-level identification—an underlying assumption of some expertise-based models [50]. Rather, we show here that the invariance hypothesis implies an algorithmic role that could be fulfilled by the mere clustering of selectivity. Consistent with the idea of a canonical cortical microcircuit [51, 52], the computations performed in each subsystem may be quite similar to the computations performed in the others. To a first approximation, the only difference between ventral stream modules could be the object category for which they are responsible.

## 2 Results

### 2.1 Theory sketch

To make the invariance hypothesis precise, let *g*_*θ*_ denote a transformation with parameter *θ*. Two images *I, I*^*′*^ depict the same object whenever *∃θ*, such that *I*^*′*^ = *g*_*θ*_*I*. For a small positive constant *ϵ*, the invariance hypothesis is the claim that the computational goal of the ventral stream is to compute a function *μ*, called a *signature*, such that

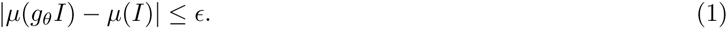

We say that a signature for which (1) is satisfied (for all *θ*) is *E*-invariant to the family of transformations *{g_θ_}*. An *E*-invariant signature that is unique to an object can be used to discriminate images of that object from images of other objects. In the context of a hierarchical model of the ventral stream, the “top level” representation of an image is its signature.

One approach to modeling the ventral stream, first taken by Fukushima’s Neocognitron [53], and followed by many other models [54, 55, 56, 57, 58], is based on iterating a basic module inspired by Hubel and Wiesel’s proposal for the connectivity of V1 simple (AND-like) and complex (OR-like) cells. In the case of HMAX [55], each “HW”-module consists of one C-unit (corresponding to a complex cell) and all its afferent S-units (corresponding to simple cells); see fig. 1-B. The response of an S-unit to an image *I* is typically modeled by a dot product with a stored template *t*, indicated here by ⟪*I, t*⟫. Since ⟪*I, t*⟫ is maximal when *I* = *t* (assuming that *I* and *t* have unit norm), we can think of an S-unit’s response as a measure of *I*’s similarity to *t*. The module corresponding to Hubel and Wiesel’s original proposal had several S-units, each detecting their stored template at a different position. Let 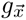 be the translation operator: when applied to an image, 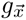 returns its translation by 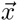. This lets us write the response of the specific S-unit which signals the presence of template *t* at position 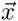 as 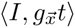. Then, introducing a nonlinear *pooling function*, which for HMAX would be the max function, the response *C*(*I*) of the C-unit (equivalently: the output of the HW-module, one element of the signature) is given by

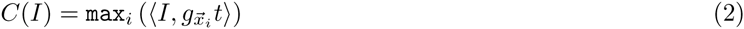

where the `max` is taken over all the S-units in the module. The region of space covered by a module’s S-units is called its *pooling domain* and the C-unit is said to pool the responses of its afferent S-units. HMAX, as well as more recent models based on this approach typically also pool over a range of scales [56, 57, 58]. In most cases, the first layer pooling domains are small intervals of translation and scaling. In the highest layers the pooling domains are usually global, i.e. over the entire range of translation and scaling that is visible during a single fixation. Notice also that this formulation is more general than HMAX. It applies to a wide class of hierarchical models of cortical computation, e.g., [53, 59, 60, 58]. For instance, *t* need not be directly interpretable as a template depicting an image of a certain object. A convolutional neural network in the sense of [61, 62] is obtained by choosing *t* to be a “prototype” obtained as the outcome of a gradient descent-based optimization procedure. In what follows we use the HW-module language since it is convenient for stating the domain-specificity argument.

HW-modules can compute approximately invariant representations for a broad class of transformations [37]. However, and this is a key fact: the conditions that must be met are different for different transformations. Following Anselmi et al. [37], we can distinguish two “regimes”. The first regime applies to the important special case of transformations with a group structure, e.g., 2D affine transformations. The second regime applies more broadly to any locally-affine transformation.

For a family of transformations *{g_θ_}*, define the *orbit* of an image *I* to be the set *{O*_*I*_ = *g*_*θ*_*I, θ* ∈ ℝ *}*. Anselmi et al. [37] proved that HW-modules can pool over other transformations besides translation and scaling. It is possible to pool over any transformation for which orbits of template objects are available. A biologically-plausible way to learn the pooling connections within an HW-module could be to associate temporally adjacent frames of the video of visual experience (as in e.g., [63, 64, 65, 66, 67, 68]). In both regimes, the following condition is required for the invariance obtained from the orbits of a set of template objects to generalize to new objects. For all *g*_*θ*_*I ∈ O*_*I*_ there is a corresponding *g*_*θ*_ *t* ∈ *O*_*t*_ such that

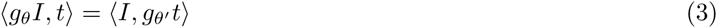

In the first regime, eq. (3) holds regardless of the level of similarity between the templates and test objects. Almost any templates can be used to recognize any other images invariantly to group transformations (see SI Text 1). Note also that this is consistent with reports in the literature of strong performance achieved using random filters in convolutional neural networks [69, 70, 71]. Figure 1-A illustrates that the orbit with respect to in-plane rotation is invariant.

In the second regime, corresponding to non-group transformations, it is not possible to achieve a perfect invariance. These transformations often depend on information that is not available in a single image. For example, rotation in depth depends on an object’s 3D structure and illumination changes depend on its material properties (see Text S1). Despite this, approximate invariance to smooth non-group transformations can still be achieved using prior knowledge of how similar objects transform. Second-regime transformations are *class-specific*, e.g., the transformation of object appearance caused by a rotation in depth is not the same 2D transformation for two objects with different 3D structures. However, by restricting to a class where all the objects have similar 3D structure, all objects do rotate (approximately) the same way. Moreover, this commonality can be exploited to transfer the invariance learned from experience with (orbits of) template objects to novel objects seen only from a single example view.

### 2.2 Simulations: Core predictions

The theory makes two core predictions:

1. Learned invariance to group transformations should be transferable from any set of stimuli to any other.
2. For non-group transformations, approximate invariance will transfer within certain object classes. In the case of 3D depth-rotation, it will transfer within classes for which all members share a common 3D structure.

Both core predictions were addressed with tests of transformation-tolerant recognition based on a single example view. Two image sets were created to test the first core prediction: (A) 100 faces derived from the Max-Planck institute face dataset [72]. Each face was oval-cropped to remove external features and normalized so that all images had the same mean and variance over pixels (as in [73]). (B) 100 random greyscale noise patterns. 29 images of each face and random noise pattern were created by placing the object over the horizontal interval from 40 pixels to the left of the image’s center up to 40 pixels to the right of the image’s center in increments of 5 pixels. All images were 256× 256 pixels.

Three image sets were created to test the second core prediction: (A) 40 untextured face models were rendered at each orientation in 5*°* increments from 95*°* to 95*°*. (B) 20 objects sharing a common gross structure (a conical shape) and differing from one another by the exact placement and size of smaller bumps. (C) 20 objects sharing gross structure consisting of a central pyramid on a flat plane and two walls on either side. Individuals differed from one another by the location and slant of several additional bumps. The face models were generated using Facegen [74]. Class B and C models were generated with Blender [75]. All rendering was also done with Blender and used perspective projection at a resolution of 256× 256 pixels.

The tests of transformation-tolerant recognition from a single example were performed as follows. In each “block”, the model was shown a reference image and a set of query images. The reference image always depicted an object under the transformation with the median parameter value. That is, for rotation in depth of faces, it was a frontal face (0*°*) and for translation, the object was located in the center of the visual field. Each query image either depicted the same object as the reference image (target case) or a different object (distractor case). In each block, each query image was shown at each position or angle in the block’s testing interval. All testing intervals were symmetric about 0. Using a sequence of testing intervals ordered by inclusion, it was possible to investigate how tolerance declines with increasingly demanding transformations. The radius of the testing interval is the abscissa of the plots in figures 2 and 3.

For each repetition of the translation experiments, 30 objects were randomly sampled from the template class and 30 objects from the testing class. For each repetition of the depth-rotation experiments, 10 objects were sampled from template and testing classes that were always disjoint from one another.

Networks consisting of *K* HW-modules were constructed where *K* was the number of sampled template objects. The construction followed the procedure described in the method section below. Signatures computed by these networks are vectors with *K* elements. In each block, the signature of the reference image was compared to the signature of each query image by its Pearson correlation and ranked accordingly. This ranked representation provides a convenient way to compute the ROC curve since it admits acceptance thresholds in terms of ranks (as opposed to real numbers). Thus, the final measure of transformation tolerance reported on the ordinate of the plots in figures (2) and (3) is the mean area under the ROC curve (AUC) over all choices of reference object and repetitions of the experiment with different training / test set splits. Since AUC is computed by integrating over acceptance thresholds, it is a bias free statistic. In this case it is analogous to *d*^*′*^ for the corresponding 2AFC same-different task. When performance is invariant, AUC as a function of testing interval radius will be a flat line.

If there is imperfect invariance (*ϵ*-invariance), then performance will decline as the radius of the testing interval is increased. To assess imperfect invariance, it is necessary to compare with an appropriate baseline at whatever performance level would be achieved by similarity in the input. Since any choice of input encoding induces its own similarity metric, the most straightforward way to obtain interpretable results is to use the raw pixel representation as the baseline (red curves in figures 2 and 3). Thus, a one layer architecture was used for these simulations: each HW-module directly receives the pixel representation of the input.

The first core prediction was addressed by testing translation-tolerant recognition with models trained using random noise templates to identify faces and vice versa (figure 2). The results in the plots on the diagonal for the view-based model (blue curve) indicate that face templates can indeed be used to identify other faces invariantly to translation; and random noise templates can be used to identify random noise invariantly to translation. The key prediction of the theory concerns the off-diagonal plots. In those cases, templates from faces were used to recognize noise patterns and noise was used to recognize faces. Performance was invariant in both cases; the blue curves in figure 2 were flat. This result was in accord with the theory’s prediction for the group transformation case: the templates need not resemble the test images.

**Figure 2:**
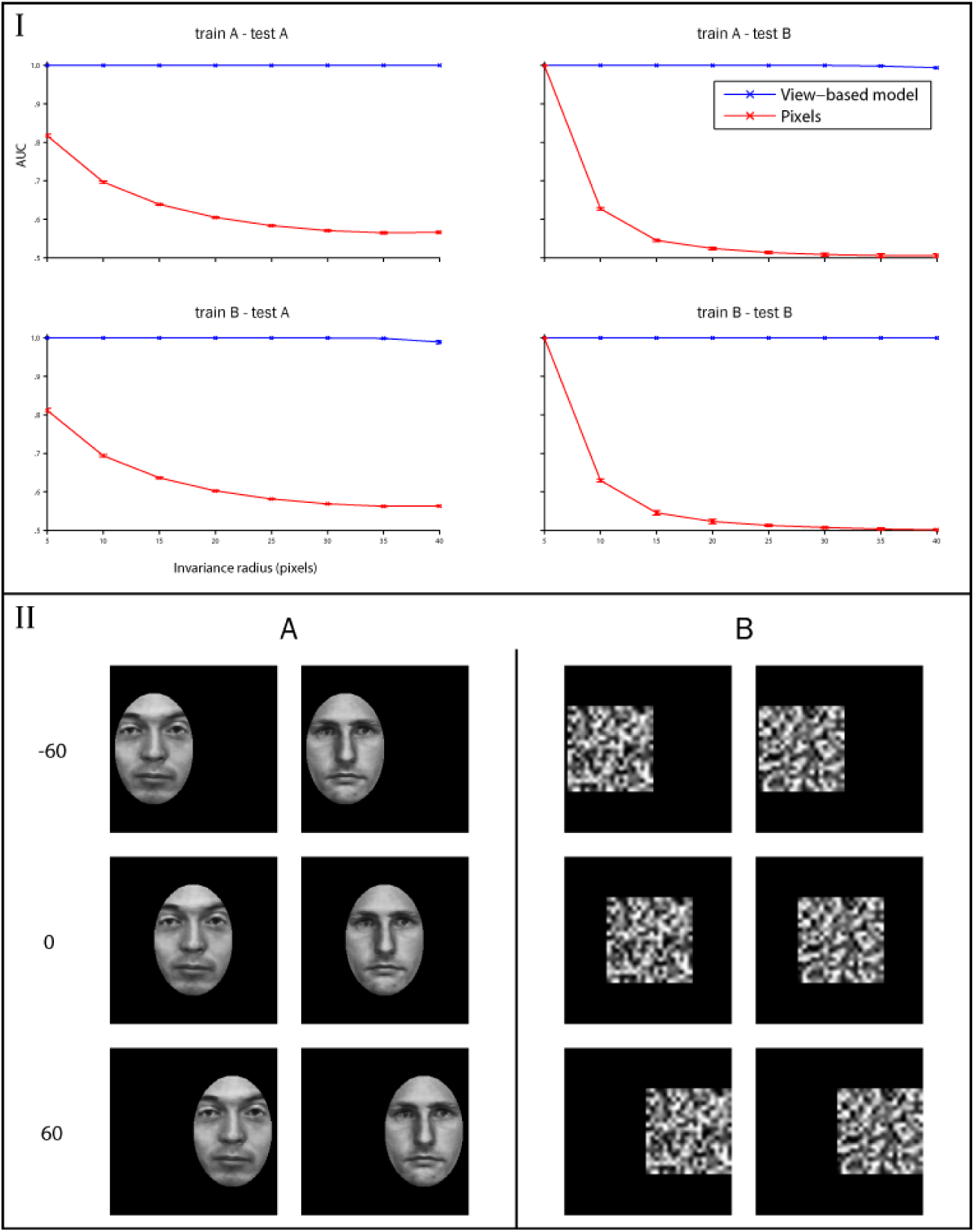
Translation invariance. Bottom panel (II): Example images from the two classes. The faces were obtained from the Max-Planck Institute dataset [72] and then contrast normalized and translated over a black background. Top panel (I): The left column shows the results of a test of translation invariance for faces and the right column shows the same test for random noise patterns. The view-based model (blue curve) was built using templates from class A in the top row and class B in the bottom row. The abscissa of each plot shows the maximum invariance range (a distance in pixels) over which target and distractor images were presented. The view-based model was never tested on any of the images that were used as templates. Error bars (*±*1 standard deviation) were computed over 5 repetitions of the experiment using different (always disjoint) sets of template and testing images.

The second core prediction concerning class-specific transfer of learned *ϵ*-invariance for non-group transformations was addressed by analogous experiments with 3D depth-rotation. Transfer of invariance both within and between classes was assessed using 3 different object classes: faces and two synthetic classes. The level of rotation tolerance achieved on this difficult task was the amount by which performance of the view-based model (blue curve) exceeded the raw pixel representation’s performance for the plots on the diagonal of figure 3. The off-diagonal plots show the deleterious effect of using templates from the wrong class.

There are many other non-group transformations besides depth-rotation. Text S1 describes additional simulations for changes in illumination. These depend on material properties. It also describes simulations of pose (standing, sitting, etc)-invariant body recognition.

**Figure 3:**
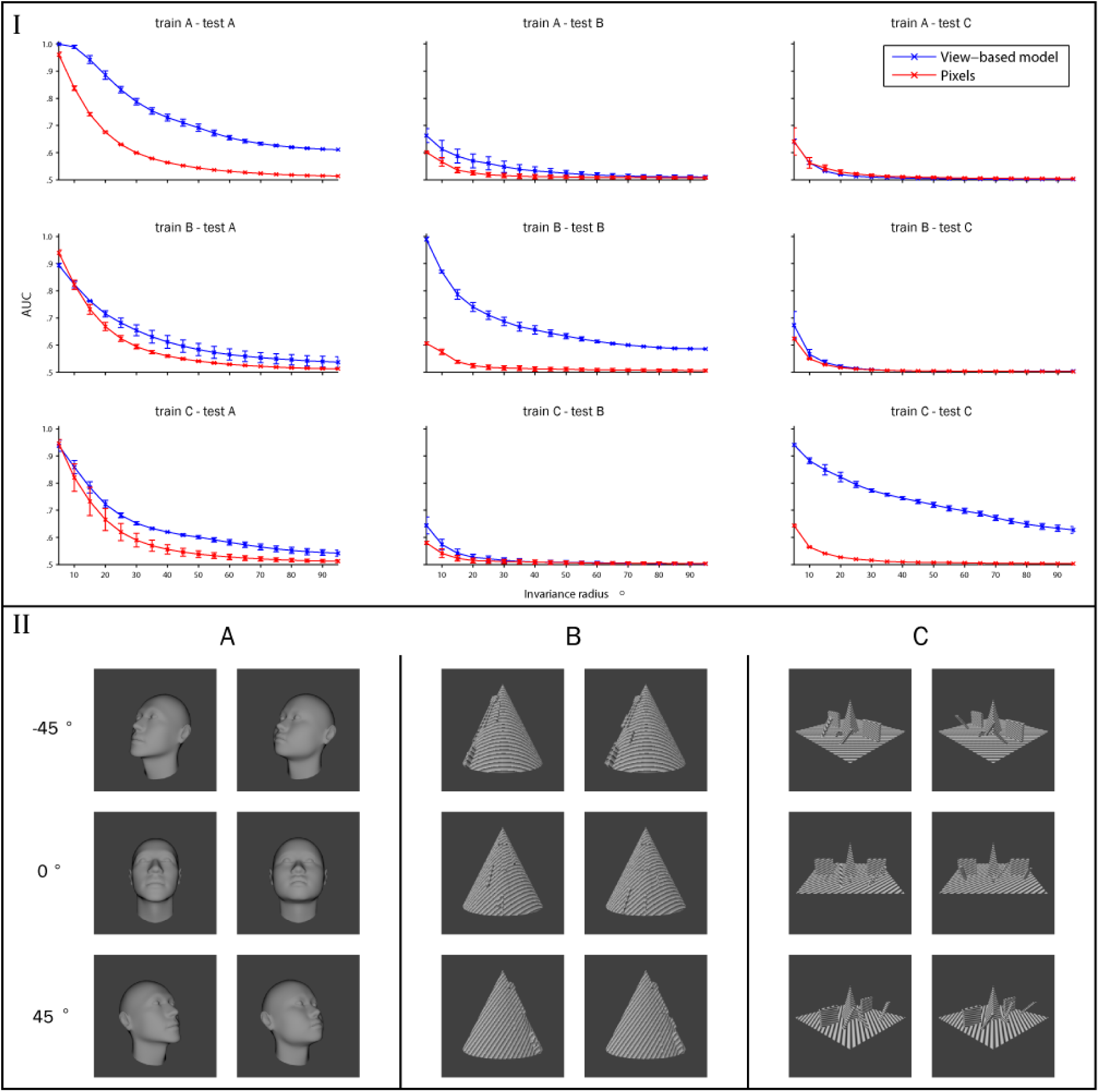
Class-specific transfer of depth-rotation invariance. Bottom panel (II): Example images from the three classes. Top panel (I): The left column shows the results of a test of 3D rotation invariance on faces (class A), the middle column shows results for class B and the right column shows the results for class C. The view-based model (blue curve) was built using images from class A in the top row, class B in the middle row, and class C in the bottom row. The abscissa of each plot shows the maximum invariance range (degrees of rotation away from the frontal face) over which target and distractor images were presented. The view-based model was never tested on any of the images that were used as templates. Error bars (*±*1 standard deviation) were computed over 20 cross validation runs using different choices of template and test images. Only the plots on the diagonal (train A - test A, train B - test B, train C - test C) show an improvement of the view-based model over the pixel representation. That is, only when the test images transform similarly to the templates is there any benefit from pooling.

### 2.3 Transformation compatibility

How can object experience—i.e., templates—be assigned to subsystems in order to facilitate productive transfer? If each individual object is assigned to a separate group, the negative effects of using templates from the wrong class are avoided; but past experience can never be transferred to new objects. So far we have only said that “3D structure” determines which objects can be productively grouped together. In this section we derive a more concrete criterion: transformation compatibility.

Given a set of objects sampled from a category, what determines when HW-modules encoding templates for a few members of the class can be used to approximately invariantly recognize unfamiliar members of the category from a single example view? Recall that the transfer of invariance depends on the condition given by eq. (3). For non-group transformations this turns out to require that the objects “transform the same way” (see Text S1 for the proof; the notion of a “nice class” is also related [76, 77]). Given a set of orbits of different objects (only the image sequences are needed), we would like to have an index 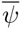 that measures how similarly the objects in the class transform. If an object category has too low 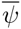, then there would be no gain from creating a subsystem for that category. Whenever a category has high 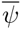, it is a candidate for having a dedicated subsystem.

The transformation compatibility of two objects *A* and *B* is defined as follows. Consider a smooth transformation *T* parameterized by *i*. Since *T* may be class-specific, let *T*_*A*_ denote its application to object *A*. One of the requirements that must be satisfied for *E*-invariance to transfer from an object *A* to an object *B* is that *T*_*A*_ and *T*_*B*_ have equal Jacobians (see Text S1). This suggests an operational definition of the transformation compatibility between two objects *Ψ*(*A, B*).

Let *A*_*i*_ be the *i*_*th*_ frame of the video of object A transforming and *B*_*i*_ be the *i*_*th*_ frame of the video of object B transforming. The Jacobian can be approximated by the “video” of difference images: *J*_*A*_(*i*) = *|A*_*i*_ - *A*_*i*+1_*|* (*∀i*). Then define the “instantaneous” transformation compatibility *Ψ*(*A, B*)(*i*) := *(J*_*A*_(*i*), *J*_*B*_(*i*)). Thus for a range of parameters *i ϵ R* = [*-r, r*], the empirical transformation compatibility between *A* and *B* is

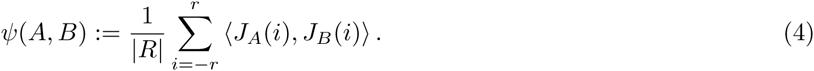

The index 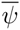 that we compute for sets of objects is the mean value of *Ψ*(*A, B*) taken over all pairs *A, B* from the set. For very large sets of objects it could be estimated by randomly sampling pairs. In the present case, we were able to use all pairs in the available data.

For the case of rotation in depth, we used 3D modeling / rendering software [75] to obtain (dense samples from) orbits. We computed the transformation compatibility index 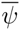 for several datasets from different sources. Faces had the highest 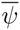 of any naturalistic category we tested—unsurprising since recognizability likely influenced face evolution. A set of chair objects (from [78]) had very low 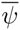 implying no benefit would be obtained from a chair-specific region. More interestingly, we tested a set of synthetic “wire” objects, very similar to those used in many classic experiments on view-based recognition e.g. [79, 80, 81]. We found that the wire objects had the lowest 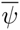 of any category we tested; experience with familiar wire objects does not transfer to new wire objects. Therefore it is never productive to group them into a subsystem.

**Table 1:** Table of transformation compatibilities. COIL-100 is a library of images of 100 common household items photographed from a range of orientations using a turntable [82]. The wire objects resemble those used in psychophysics and physiology experiments: [79, 80, 81]. They were generated according to the same protocol as in those studies.

| Object class | Transformation | $\bar{\psi}$ |
| --- | --- | --- |
| Chairs | Rotation in depth | 0.00540 |
| Fig. 3 faces | Rotation in depth | 0.57600 |
| Fig. 3 class B | Rotation in depth | 0.95310 |
| Fig. 3 class C | Rotation in depth | 0.83800 |
| Fig. 3 all classes | Rotation in depth | 0.26520 |
| COIL-100 [82] | Rotation in depth | 0.00630 |
| Wire objects [80] | Rotation in depth | -0.00007 |

### 2.4 Simulations: The domain specific architecture of visual cortex

The above considerations suggest an unsupervised strategy for sorting object experience into subsystems. An online *Ψ*-based clustering algorithm could sort each newly learned object representation into the subsystem (cluster) with which it transforms most compatibly. With some extra assumptions beyond those required for the main theory, such an algorithm could be regarded as a very stylized model of the development (or evolution) of visual cortex. In this context we asked: Is it possible to derive predictions for the specific object classes that will “get their own private piece of real estate in the brain” [8] from the invariance hypothesis?

The extra assumptions required at this point are as follows.

1. Cortical object representations (HW-modules) are sampled from the distribution *D* of objects and their transformations encountered under natural visual experience.
2. Subsystems are localized on cortex.
3. The number of HW-modules in a local region and the proportion belonging to different categories determines the predicted BOLD response for contrasts between the categories. For example, a cluster with 90% face HW-modules, 10% car HW-modules, and no other HW-modules would respond strongly in the faces - cars contrast, but not as strongly as it would in a faces - airplanes contrast. We assume that clusters containing very few HW-modules are too small to be imaged with the resolution of fMRI—though they may be visible with other methods that have higher resolution.

Any model that can predict which specific categories will have domain-specific regions must depend on contingent facts about the world, in particular, the—difficult to approximate—distribution *D* of objects and their transformations encountered during natural vision. Consider the following: HW-modules may be assigned to cluster near one another on cortex in order to maximize the transformation compatibility 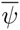 of the set of objects represented in each local neighborhood. Whenever a new object is learned, its HW-module could be placed on cortex in the neighborhood with which it transforms most compatibly. Assume a new object is sampled from *D* at each iteration. We conjecture that the resulting cortex model obtained after running this for some time would have a small number of very large clusters, probably corresponding to faces, bodies, and orthography in a literate brain’s native language. The rest of the objects would be encoded by HW-modules at random locations. Since neuroimaging methods like fMRI have limited resolution, only the largest clusters would be visible to them. Cortical regions with low 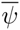 would appear in neuroimaging experiments as generic “object regions” like LOC [83].

Since we did not attempt the difficult task of sampling from *D*, we were not able to test the conjecture directly. However, by assuming particular distributions and sampling from a large library of 3D models [78, 74], we can study the special case where the only transformation is rotation in depth. Each object was rendered at a range of viewpoints: 90*°* to 90*°* in increments of 5 degrees. The objects were drawn from five categories: faces, bodies, animals, chairs, and vehicles. Rather than trying to estimate the frequencies with which these objects occur in natural vision, we instead aimed for predictions that could be shown to be robust over a range of assumptions on *D* Thus we repeated the online clustering experiment three times, each using a different object distribution (see Table S2, and Figures S6, S7, S8, S9, and S10).

The *Ψ*-based clustering algorithm we used can be summarized as follows: Consider a model consisting of a number of subsystems. When an object is learned, add its newly-created HW-module to the subsystem with which its transformations are most compatible. If the new object’s average compatibility with all the existing subsystems is below a threshold, then create a new subsystem for the newly learned object. Repeat this procedure for each object—sampled according to the distribution of objects encountered in natural vision (or whatever approximation is available). See Text S1 for the algorithm’s pseudocode. Figure 4 shows example clusters obtained by this method.

**Figure 4:**
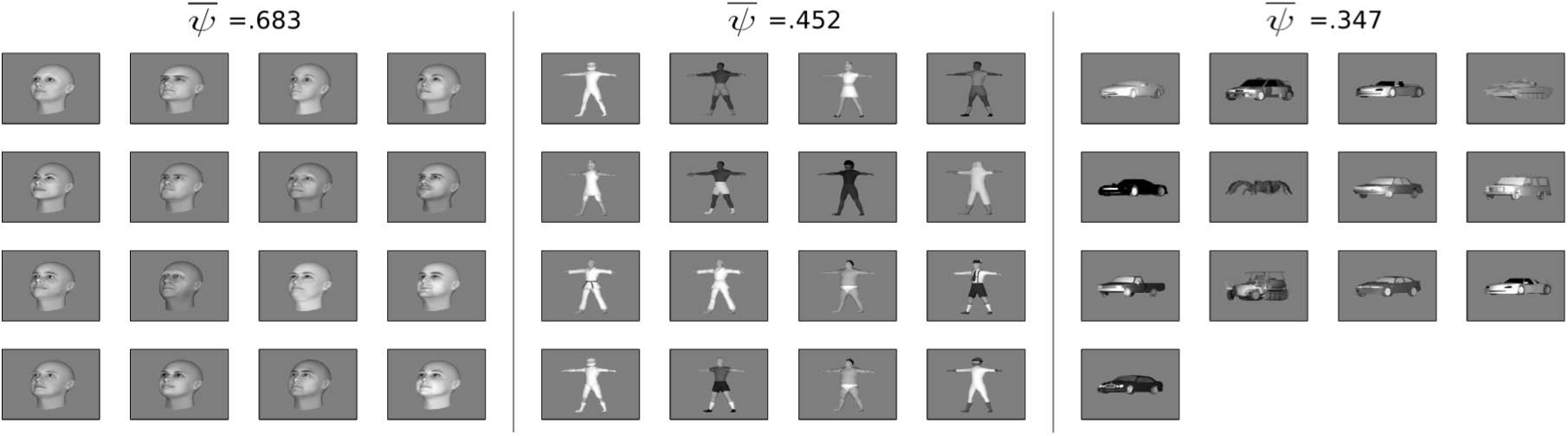
Example clustering results. Three example clusters that developed in a simulation with an object distribution biased against faces (the same simulation as in figures S8-C, S9-C, and S10-C).

**Figure 5:**
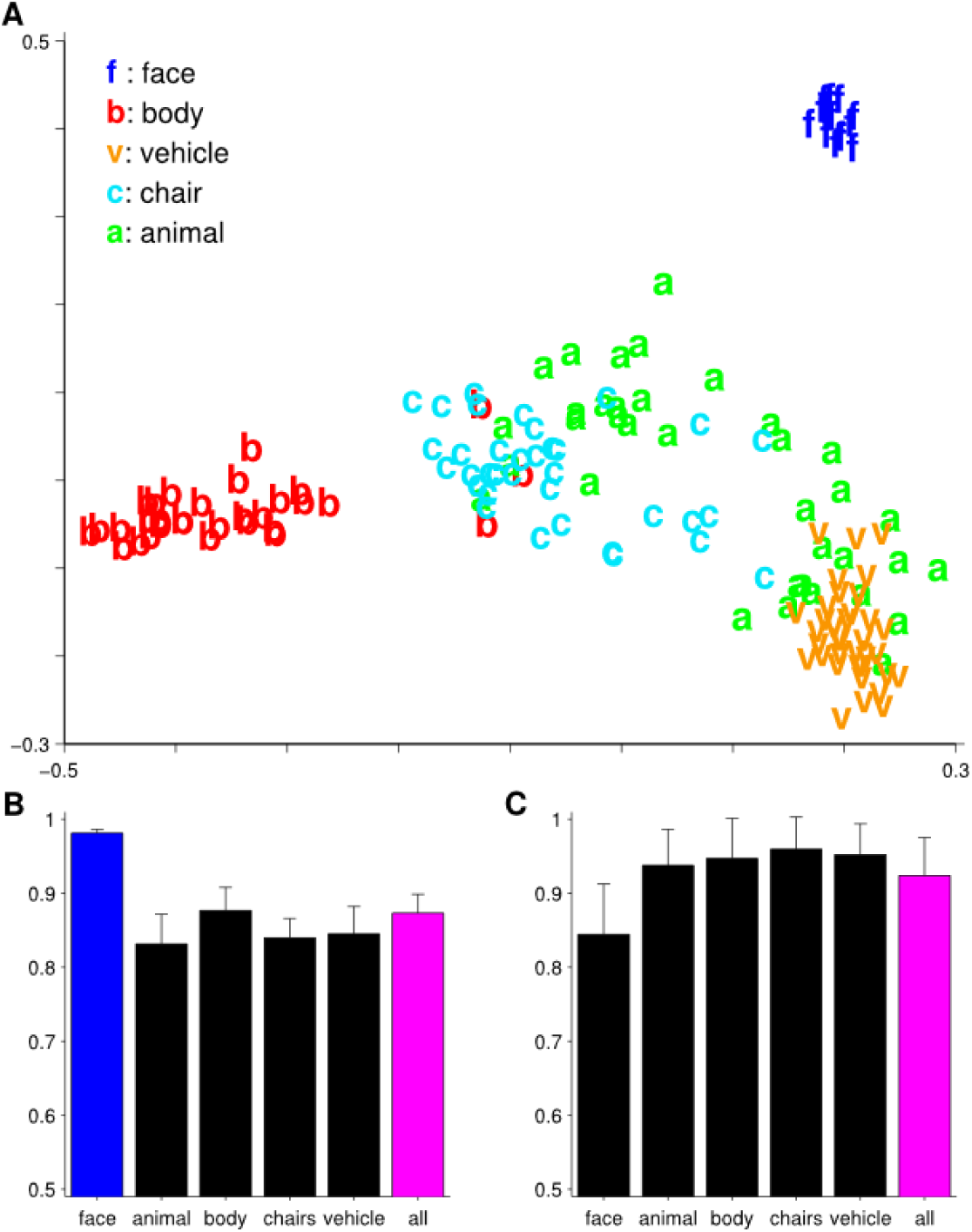
Simulation of the development of domain-specific regions. In this case the distribution of objects was biased against faces (faces were only 16 of the 156 objects in this simulation). Depth-rotation is the only transformation used here. The main assumption is that the distance along cortex between two HW-modules for two different templates is proportional to how similarly the two templates transform. See Figures S8, S9, and S10 for results of the analogous simulations using different object distributions **A.** Multidimensional scaling plot based on pairwise transformation compatibility *Ψ*. **B.** Results on a test of view-invariant face verification (same-different matching). Each bar corresponds to a different cluster produced by an iterative clustering algorithm based on 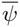 which models visual development—see supplementary methods. The labels on the abscissa correspond to the dominant category in the cluster. **C.** Basic-level categorization results: Cars versus airplanes. Error bars were obtained by repeating the experiment 5 times, presenting the objects in a different random order during development and randomly choosing different objects for the test set.

Robust face and body clusters always appeared (figure 5, SI figures S8, S9, and S10). Due to the strong effect of *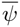*, a face cluster formed even when the distribution of objects was biased *against* faces as in figure 5. Most of the other objects ended up in very small clusters consisting of just a few objects. For the experiment of figures 4 and 5, 16% of the bodies, 64% of the animals, 44% of the chairs, and 22% of the vehicles were in clusters consisting of just one object. No faces ended up in single-object clusters.

To confirm that *Ψ*-based clustering is useful for object recognition with these images, we compared the recognition performance of the subsystems to the complete system that was trained using all available templates irrespective of their cluster assignment. We simulated two recognition tasks: one basic-level categorization task, view-invariant cars vs. airplanes, and one subordinate-level task, view-invariant face recognition. For these tests, each “trial” consisted of a pair of images. In the face recognition task, the goal was to respond ‘same’ if the two images depicted the same individual. In the cars vs. airplanes case, the goal was to respond ‘same’ if both images depicted objects of the same category. In both cases, all the objects in the cluster were used as templates; the test sets were completely disjoint. The classifier was the same as in figures 2 and 3. In this case, the threshold was optimized on a held out training set.

As expected from the theory, performance on the subordinate-level view-invariant face recognition task was significantly higher when the face cluster was used (figure 5-B). The basic-level categorization task was performed to similar accuracy using any of the clusters (figure 5-C). This confirms that invariance to class-specific transformations is only necessary for subordinate level tasks.

## Discussion

We explored implications of the hypothesis that achieving transformation invariance is the main goal of the ventral stream. Invariance from a single example could be achieved for group transformations in a generic way. However, for non-group transformations, only approximate invariance is possible; and even for that, it is necessary to have experience with objects that transform similarly. This implies that the optimal organization of the ventral stream is one that facilitates the transfer of invariance within—but not between—object categories. Assuming that a subsystem must reside in a localized cortical neighborhood, this could explain the function of domain-specific regions in the ventral stream’s recognition algorithm: to enable subordinate level identification of novel objects from a single example.

Following on from our analysis implicating transformation compatibility as the key factor determining when invariance can be productively transferred between objects, we simulated the development of visual cortex using a clustering algorithm based on transformation compatibility. This allowed us to address the question of why faces, bodies, and words get their own dedicated regions but other object categories (apparently) do not [8]. This question has not previously been the focus of theoretical study.

Despite the simplicity of our model, we showed that it robustly yields face and body clusters across a range of object frequency assumptions. We also used the model to confirm two theoretical predictions: (1) that invariance to non-group transformations is only needed for subordinate level identification; and (2) that clustering by transformation compatibility yields subsystems that improve performance beyond that of the system trained using data from all categories. These results motivate the the next phase of this work: building biologically-plausible models that learn from natural video. Such models automatically incorporate a better estimate of the natural object distribution. Variants of these models may be able to quantitatively reproduce human level performance on simultaneous multi-category subordinate level (i.e., fine-grained) visual recognition tasks and potentially find application in computer vision as well as neuroscience. In [42], we report encouraging preliminary results along these lines.

Why are there domain-specific regions in later stages of the ventral stream hierarchy but not in early visual areas [2, 3]? The templates used to implement invariance to group transformations need not be changed for different object classes while the templates implementing non-group invariance are class-specific. Thus it is efficient to put the generic circuitry of the first regime in the hierarchy’s early stages, postponing the need to branch to different domain-specific regions tuned to specific object classes until later, i.e., more anterior, stages. In the macaque face-processing system, category selectivity develops in a series of steps; posterior face regions are less face selective than anterior ones [34, 84]. Additionally, there is a progression from a view-specific face representation in earlier regions to a view-tolerant representation in the most anterior region [34]. Both findings could be accounted for in a face-specific hierarchical model that increases in template size and pooling region size with each subsequent layer (e.g., [85, 86, 41, 42]). The use of large face-specific templates may be an effective way to gate the entrance to the face-specific subsystem so as to keep out spurious activations from non-faces. The algorithmic effect of large face-specific templates is to confer tolerance to clutter [41, 42]. These results are particularly interesting in light of models showing that large face templates are sufficient to explain holistic effects observed in psychophysics experiments [87, 73].

As stated in the introduction, properties of the ventral stream are thought to be determined by three factors: computational and algorithmic constraints; (2) biological implementation constraints; and (3) the contingencies of the visual environment [18, 19, 20, 21, 22]. Up to now, we have stressed the contribution of factor (1) over the others. In particular, we have almost entirely ignored factor (2). We now discuss the role played by anatomical considerations in this account of ventral stream function. That the the circuitry comprising a subsystem must be localized on cortex is a key assumption of this work. In principle, any HW-module could be anywhere, as long as the wiring all went to the right place. However, there are several reasons to think that the actual constraints under which the brain operates and its available information processing mechanisms favor a situation in which, at each level of the hierarchy, all the specialized circuitry for one domain is in a localized region of cortex, separate from the circuitry for other domains. Wiring length considerations are likely to play a role here [88, 89, 90, 91]. Another possibility is that localization on cortex enables the use of neuromodulatory mechanisms that act on local neighborhoods of cortex to affect all the circuitry for a particular domain at once [92].

There are other domain-specific regions in the ventral stream besides faces and bodies; we consider several of them in light of our results here. It is possible that even more regions for less-common (or less transformation-compatible) object classes would appear with higher resolution scans. One example may be the fruit area, discovered in macaques with high-field fMRI [3].

1. **Lateral Occipital Complex (LOC) [83]** These results imply that LOC is not really a dedicated region for general object processing. Rather, it is a heterogeneous area of cortex containing many domain-specific regions too small to be detected with the resolution of fMRI. It may also include clusters that are not dominated by one object category as we sometimes observed appearing in simulations (see figure 4 and S.I. Text 1).
2. **The Visual Word Form Area (VWFA) [4]** In addition to the generic transformations that apply to all objects, printed words undergo several non-generic transformations that never occur with other objects. We can read despite the large image changes occurring when a page is viewed from a different angle. Additionally, many properties of printed letters change with typeface, but our ability to read—even in novel fonts—is preserved. Reading hand-written text poses an even more severe version of the same computational problem. Thus, VWFA is well-accounted for by the invariance hypothesis. Words are frequently-viewed stimuli which undergo class-specific transformations. This account appears to be in accord with others in the literature [93, 94].
3. **Parahippocampal Place Area (PPA) [95]** A recent study by Kornblith et al. describes properties of neurons in two macaque scene-selective regions deemed the lateral and medial place patches (LPP and MPP) [96]. While homology has not been definitively established, it seems likely that these regions are homologous to the human PPA [97]. Moreover, this scene-processing network may be analogous to the face-processing hierarchy of [34]. In particular, MPP showed weaker effects of viewpoint, depth, and objects than LPP. This is suggestive of a scene-processing hierarchy that computes a representation of scene-identity that is (approximately) invariant to those factors. Any of them might be transformations for which this region is compatible in the sense of our theory. One possibility, which we considered in preliminary work, is that invariant perception of scene identity despite changes in monocular depth signals driven by traversing a scene (e.g., linear perspective) could be discounted in the same manner as face viewpoint. It is possible that putative scene-selective categories compute depth-tolerant representations. We confirmed this for the special case of long hallways differing in the placement of objects along the walls: a view-based model that pools over images of template hallways can be used to recognize novel hallways [98]. Furthermore, fast same-different judgements of scene identity tolerate substantial changes in perspective depth [98]. Of course, this begs the question: of what use would be a depth-invariant scene representation? One possibility could be to provide a landmark representation suitable for anchoring a polar coordinate system [99]. Intriguingly, [96] found that cells in the macaque scene-selective network were particularly sensitive to the presence of long straight lines—as might be expected in an intermediate stage on the way to computing perspective invariance.

Is this proposal at odds with the literature emphasizing the view-dependence of human vision when tested on subordinate level tasks with unfamiliar examples—e.g.[79, 72, 100]? We believe it is consistent with most of this literature. We merely emphasize the substantial view-*tolerance* achieved for certain object classes, while they emphasize the lack of complete invariance. Their emphasis was appropriate in the context of earlier debates about view-invariance [101, 102, 103, 104], and before differences between the view-tolerance achieved on basic-level and subordinate-level tasks were fully appreciated [105, 106, 107].

The view-dependence observed in experiments with novel faces [72, 108] is consistent with the predictions of our theory. The 3D structure of faces does not vary wildly within the class, but there is still some significant variation. It is this variability in 3D structure within the class that is the source of the imperfect performance in our simulations. Many psychophysical experiments on viewpoint invariance were performed with synthetic “wire” objects defined entirely by their 3D structure e.g., [79, 80, 81]. We found that they were by far, the least transformation-compatible (lowest 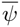) objects we tested (fig. 1). Thus our proposal predicts particularly weak performance on viewpoint-tolerance tasks with novel examples of these stimuli and that is precisely what is observed [80].

Tarr and Gauthier (1998) found that learned viewpoint-dependent mechanisms could generalize across members of a homogenous object class [107]. They tested both homogenous block-like objects, and several other classes of more complex novel shapes. They concluded that this kind of generalization was restricted to visually similar objects. These results seem to be consistent with our proposal. Additionally, our hypothesis predicts better within-class generalization for object classes with higher 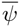. That is, transformation compatibility, not visual similarity per se, may be the factor influencing the extent of within-class generalization of learned view-tolerance. Though, in practice, the two are usually correlated and hard to disentangle. In a related experiment, Sinha and Poggio (1996) showed that the perception of an ambiguous transformation’s rigidity could be biased by experience [109]. View-based accounts of their results predict that the effect would generalize to novel objects of the same class. Since this effect can be obtained with particularly simple stimuli, it might be possible to design them so as to separate specific notions of visual similarity and transformation compatibility. In accord with our prediction that group transformations ought to be discounted earlier in the recognition process, [109] found that their effect was spared by presenting the training and test objects at different scales.

Many authors have argued that seemingly domain-specific regions are actually explained by perceptual expertise [110, 24, 26, 25, 27]. Our account is compatible with some aspects of this idea. However, it is largely agnostic about whether the sorting of object classes into subsystems takes place over the course of evolution or during an organism’s lifetime. A combination of both is also possible—e.g. as in [111]. That said, our proposal does intersect this debate in several ways.

1. Our theory agrees with most expertise-based accounts that subordinate-level identification is the relevant task.
2. The expertise argument has always relied quite heavily on the idea that discriminating individuals from similar distractors is somehow difficult. Our account allows greater precision: the precise component of difficulty that matters is invariance to non-group transformations.
3. Our theory predicts a critical factor determining which objects could be productively grouped into a module that is clearly formulated and operationalized: the transformation compatibility 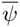.

Under our account, domain-specific regions arise because they are needed in order to facilitate the generalization of learned transformation invariance to novel category-members. Most studies of clustering and perceptual expertise do not use this task. However, Srihasam et al. tested a version of the perceptual expertise hypothesis that could be understood in this way [112]. They trained macaques to associate reward amounts with letters and numerals (26 symbols). In each trial, a pair of symbols were displayed and the task was to pick the symbol associated with greater reward. Importantly, the 3-year training process occurred in the animal’s home cage and eye tracking was not used. Thus, the distance and angle with which the monkey subjects viewed the stimuli was not tightly controlled during training. The symbols would have projected onto their retina in many different ways. These are exactly the same transformations that we proposed are the reason for the VWFA. In accord with our prediction, Srihasam et al. found that this training experience caused the formation of category-selective regions in the temporal lobe. Furthermore, the same regions were activated selectively irrespective of stimulus size, position, and font. Interestingly, this result only held for juvenile macaques, implying there may be a critical period for cluster formation[112].

Our main prediction is the link between transformation compatibility and domain-specific clustering. Thus one way to test whether this account of expertise-related clustering is correct could be to train monkeys to recognize individual objects of unfamiliar classes invariantly to 3D rotation in depth. The task should involve generalization from a single example view of a novel exemplar. The training procedure should involve exposure to videos of a large number of objects from each category undergoing rotations in depth. Several categories with different transformation compatibilities should be used. The prediction is that after training there will be greater clustering of selectivity for the classes with greater average transformation compatibility (higher *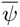*). Furthermore, if one could record from neurons in the category-selective clusters, the theory would predict some similar properties to the macaque face-processing hierarchy: several interconnected regions progressing from view-specificity in the earlier regions to view-tolerance in the later regions. However, unless the novel object classes actually transform like faces, the clusters produced by expertise should be parallel to the face clusters but separate from them.

How should these results be understood in light of recent reports of very strong performance of “deep learning” computer vision systems employing apparently generic circuitry for object recognition tasks e.g., [62, 113]? We think that exhaustive greedy optimization of parameters (weights) over a large labeled data set may have found a network similar to the architecture we describe since all the basic structural elements (neurons with nonlinearities, pooling, dot products, layers) required by our theory are identical to the elements in deep learning networks. If this were true, our theory would also explain what these networks do and why they work.

## 4 Methods

### Training HW-architectures

An *HW-architecture* refers to a feedforward hierarchical network of *HW-layers*. An HW-layer consists of *K HW-modules* arranged in parallel to one another (see Figure 1B). For an input image *I*, the output of an HW-layer is a vector *μ*(*I*) with *K* elements. If *I* depicts a particular object, then *μ*(*I*) is said to be the *signature* of that object.

The parameters (weights) of the *k*-th HW-module are uniquely determined by its *template book*

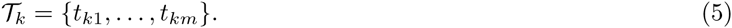

For all simulations in this paper, the output of the *k*-th HW-module is given by

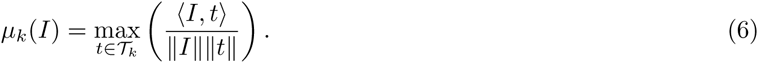

We used a nonparametric method of training HW-modules that models the outcome of temporal continuity-based unsupervised learning [67, 42]. In each experiment, the training data consisted of *K* videos represented as sequences of frames. Each video depicted the transformation of just one object. Let *G*_0_ be a family of transformations, e.g., a subset of the group of translations or rotations. The set of frames in the *k*-th video was *O*_*t*_*k* = *{gt*_*k*_ *| g ∈ G*_0_*}*.

In each simulation, an HW-layer consisting of *K* HW-modules was constructed. The template book *T*_*k*_ of the *k*-th HW-module was chosen to be

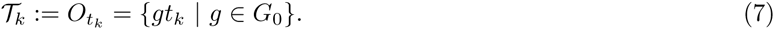

Note that HW-architectures are usually trained in a layer-wise manner (e.g., [57]). That is, layer *l* templates are encoded as “neural images” using the outputs of layer *l* − 1. However, in this paper, all the simulations use a single HW-layer.

One-layer HW-architectures are a particularly stylized abstraction of the ventral stream hierarchy. With our training procedure, they have no free parameters at all. This makes them ideal for simulations in which the aim is not to quantitatively reproduce experimental phenomena, but rather to study general principles of cortical computation that constrain all levels of the hierarchy alike.

### Experiment 1 and 2: the test of transformation-tolerance from a single example view)

**Procedure**

The training set consisted of transformation sequences of *K* template objects. At test time, in each trial the reference image was presented at the 0 transformation parameter (either 0*°*, or the center of the image for experiment 1 and 2 respectively). In each trial, a number of query images were presented, 50% of which were targets. The signature of the reference image was computed and its Pearson correlation compared with each query image. This allowed the plotting of an ROC curve by varying the acceptance threshold. The statistic reported on the ordinate of figures 2 and 3 was the area under the ROC curve averaged over all choices of reference image and all resampled training and testing sets.

#### 1. Translation experiments (Figure 2)

##### Stimuli

There were 100 faces and 100 random noise patterns in the dataset. For each repetition of the experiment, two disjoint sets of 30 objects were selected at random from the 100. The first was used as the template set and the second was used as the test set. Each experiment was repeated 5 times with different random choices of template and testing sets. The error bars on the ordinate of figure 2 are *±*1 standard deviation computed over the 5 repetitions.

#### 2. Rotation in depth experiments (Figure 3)

##### Stimuli

All objects were rendered with perspective projection. For rotation in depth experiments, the complete set of objects consisted of 40 untextured faces, 20 class B objects, and 20 class C objects. For each of the 20 repetitions of the experiment, 10 template objects and 10 test objects were randomly selected. The template and test sets were chosen independently and were always disjoint. Each face/object was rendered (using Blender [75]) at each orientation in 5*°* increments from *-*95*°* to 95*°*. The untextured face models were generated using Facegen [74].

### Experiment 3: Transformation Compatibility, Multidimensional Scaling and Online Clustering experiments (Figures 4 and 5)

#### Stimuli: Faces, bodies, vehicles, chairs and animals

Blender was used to render images of 3D models from two sources: 1. the Digimation archive (platinum edition), and 2. Facegen. Each object was rendered at a range of viewpoints: *-*90*°* to 90*°* in increments of 5 degrees. This procedure produced a transformation sequence for each object, i.e., a video. The full Digimation set consisted of *∼*10,000 objects. However, our simulations only used textured objects from the following categories: bodies, vehicles, chairs, and animals. For each experiment, the number of objects used from each class is listed in Table S2. A set of textured face models generated with FaceGen were added to the Digimation set. See Figure S7 for examples.

#### Procedure

Let *A*_*i*_ be the *i*_*th*_ frame of the video of object A transforming and *B*_*i*_ be the *i*_*th*_ frame of the video of object B transforming. Define a compatibility function *Ψ*(*A, B*) to quantify how similarly objects A and B transform.

First, approximate the Jacobian of a transformation sequence by the “video” of difference images: *J*_*A*_(*i*) = *| A*_*i*_- *A*_*i*+1_*|* (*∀i*).

Then define the pairwise transformation compatibility as:

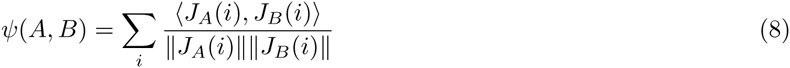

Transformation compatibility can be visualized by Multidimensional Scaling (MDS) [114]. The input to the MDS algorithm is the pairwise similarity matrix containing the transformation compatibilities between all pairs of objects.

For the *Ψ*-based online clustering experiments, consider a model consisting of a number of subsystems (HW-architectures). The clustering procedure was as follows: At each step a new object is learned. Its newly-created HW-module is added to the subsystem with which its transformations are most compatible. If the new object’s average compatibility with all the existing subsystems is below a threshold, then create a new subsystem for the newly learned object. Repeat this procedure for each object.

The objects for this experiment were sampled from three different distributions: “realistic” distribution, uniform distribution, and the biased against faces distribution, see Table S2 for the numbers of objects used from each class.

The algorithm’s pseudocode is in Text S1 (Section 5.3). Figure 4 shows examples of clusters obtained by this method.

### Experiment 4: evaluating the clustered models on subordinate-level and basic-level tasks (Figure 5)

#### Stimuli

The stimuli were the same as in experiment 3.

#### Procedure

To confirm that *Ψ*-based clustering is useful for object recognition with these images, we compared the recognition performance of the subsystems to the complete system that was trained using all available templates irrespective of their subsystem assignment.

Two recognition tasks were simulated: one basic level categorization task, view-invariant cars vs. airplanes, and one subordinate level task, view-invariant face recognition. For the subordinate face recognition task, a pair of face images were given, the task was to determine whether they depict the same person (positive) or not (negative). For basic level categorization, a pair of car/airplane images were given; the task was to determine whether they depicted the same basic-level category or not. That is, whether two images are both cars (positive), both airplanes (positive) or one airplane and one car (negative). The classifier used for both tasks was the same as the one used for experiments 1 and 2: for each test pair, the Pearson correlation between the two signatures was compared to a threshold. The threshold was optimized on a disjoint training set.

For each cluster, an HW-architecture was trained using only the objects in that cluster. If there were *K* objects in the cluster, then its HW-architecture had *K* HW-modules. Applying eq. (7), each HW-module’s template book was the set of frames from the transformation video of one of the objects in the cluster. For both tasks, in the test phase, the signature of each test image was computed with eq. (6).

Since the clustering procedure depends on the order in which the objects were presented, for each of the 3 object distributions, we repeated the basic-level and subordinate level recognition tasks 5 times using different random presentation orders. The error bars in figure 5-B, 5-C, and S15 convey the variability (one standard deviation) arising from presentation order.

#### Evaluation parameters

- 60 new face objects (disjoint from the clustering set)
- Data was evenly split to 5 folds, 12 objects per fold.
- For each fold, 48 objects were used for threshold optimization. For the face recognition case, 12 faces were used for testing. For the basic-level case, 12 objects of each category were used for testing.
- For each fold, 4000 pairs were used to learn the classification threshold *θ* (see below), 4000 pairs for testing.
- Performance was averaged over all folds.

## Supplementary Information

### 5 Remarks on the theory of architectures for invariant recognition

**Figure 6:**
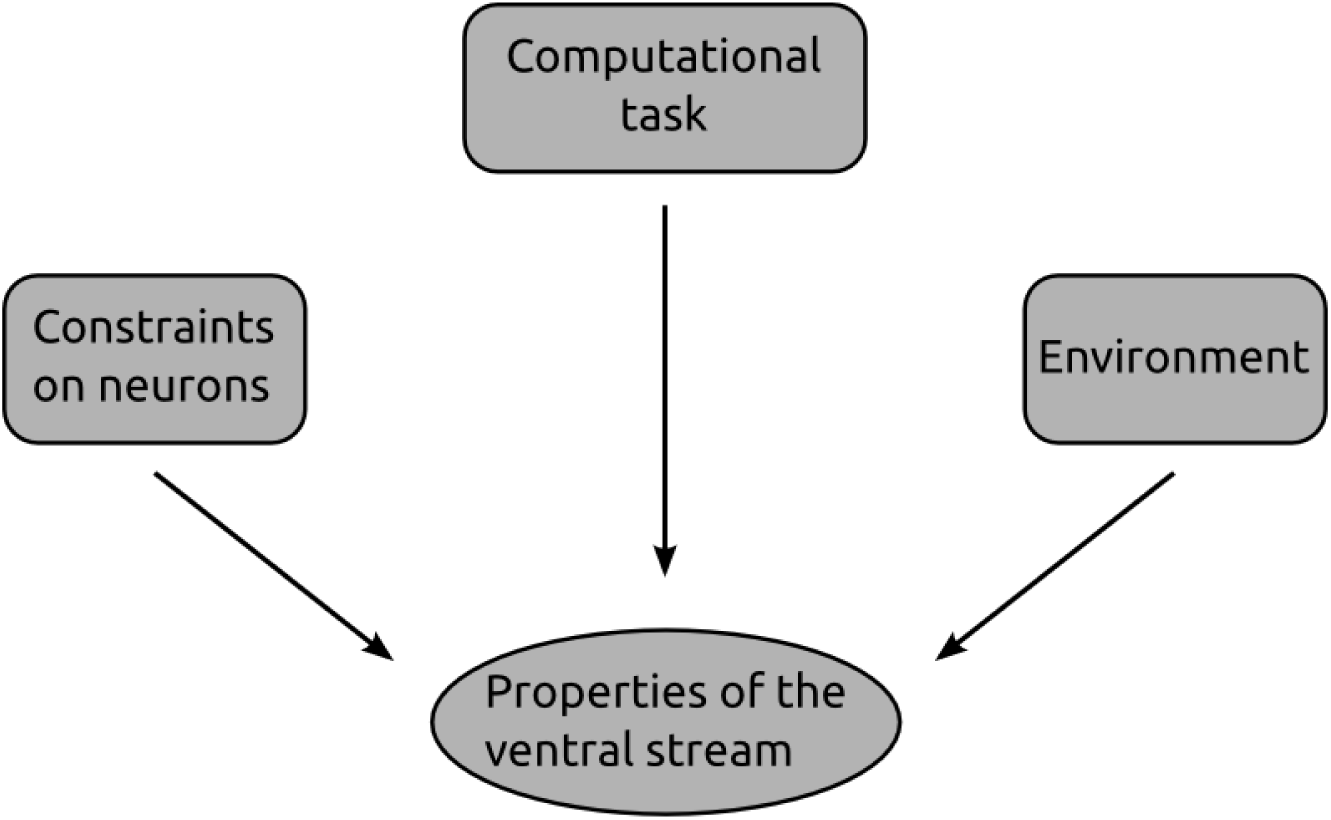
It is hypothesized that properties of the ventral stream are determined by these three factors. We are not the only ones to identify them in this way. For example, Simoncelli and Olshausen distinguished the same three factors [20]. The crucial difference between their *efficient coding hypothesis* and our *invariance hypothesis* is the particular computational task that we consider. In their case, the task is to provide an efficient representation of the visual world. In our case, the task is to provide an invariant signature supporting object recognition.

The new theory of architectures for object recognition [37]—applied here to the ventral stream—is quite general. It encompasses many non-biological hierarchical networks in the computer vision literature in addition to ventral stream models like HMAX. It also implies the existence of a wider class of hierarchical recognition algorithms that has not yet been fully explored. The conjecture with which this paper is concerned is that the algorithm implemented by the ventral stream’s feedforward processing is in this class. The theory can be developed from four postulates: (1) Computing a representation that is unique to each object and invariant to identity-preserving transformations is the main computational problem to be solved by an object recognition system—i.e., by the ventral stream. (2) The ventral stream’s feedforward, hierarchical operating mode is sufficient for recognition [115, 116, 117]. (3) Neurons can compute high-dimensional dot products between their inputs and a stored vector of synaptic weights [118]. (4) Each layer of the hierarchy implements the same basic “HW-”module, performing filtering and pooling operations via the scheme proposed by Hubel and Wiesel for the wiring of V1 simple cells to complex cells [119].

We argue that as long as these postulates are approximately correct, then the algorithm implemented by the (feedforward) ventral stream is in the class described by the theory, and this is sufficient to explain its domain-specific organization.

#### 5.1 The first regime: generic invariance

First, consider the (compact) group of 2D in-plane rotations *G*. With some abuse of notation, we use *g* to indicate both an element of *G* and its unitary representation acting on images. The orbit of an image *I* under the action of the group is *O*_*I*_ = *{gI | g ϵG}*. The orbit is invariant and unique to the object depicted in *I*. That is, *O*_*I*_ = *O*_*I′*_ if and only if *I*^*′*^ = *gI* for some *gϵG*. For an example, let *I* be an image. Its orbit *O*_*I*_ is the set of all images obtained by rotating *I* in plane. Now consider, *g*_90_*° I*, its rotation by 90*°*. The two orbits are clearly the same *O*_*I*_ = *O*_*g*90*° I*_ — the set of images obtained by rotating *I* is the same as the set of images obtained by rotating *g*_90_° *I*.

The fact that orbits are invariant and unique (for compact groups) suggests a recognition strategy. Simply store the orbit for each object. Then when new objects appear, check what orbit they are in. While that strategy would certainly work, it would be impossible to implement in practice. If we were restricted to cases where we had already stored the entire orbit then we would only ever be able to recognize objects that we had previously encountered under all their possible appearances. The key property that enables this approach to object recognition is the following condition. For a stored *template t*

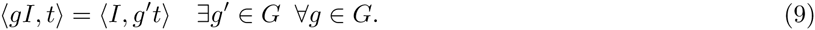

It is true whenever *g* is unitary since in that case *g*^*t*^ = *g*^−1^. It implies that it is not necessary to have the orbit of *I* before the test. Instead, the orbit of *t* is sufficient. Eq. (9) enables the invariance learned from observing a set of templates to transfer to new images. Consider the case where the full orbit of several templates *t*^1^, *…, t*^*K*^ were stored, but *I* is a completely novel image. Then an invariant signature *μ*(*·*) can be defined as

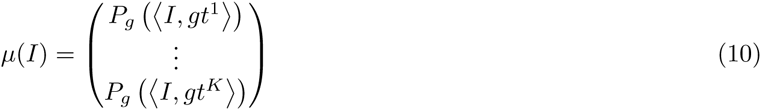

Just as in the HMAX case, *P*_*g*_ must be unchanged by permuting the order of its arguments, e.g., *P*_*g*_(·) = max_*g*_(·) or ∑_*g*_(·).

So far, this analysis has only applied to compact groups. Essentially the only interesting one is in-plane rotation. We need an additional idea in order to consider more general groups—it will also be needed later when we consider non-group transformations in the theory’s second regime. The idea is as follows. Most transformations are generally only observed through a range of transformation parameters. For example, in principle, one could translate arbitrary distances. But in practice, all translations are contained within some finite window. That is, rather than considering the full orbit under the action of *G*, we consider partial orbits under the action of a subset *G*_0_ *⊂ G* (note: *G*_0_ is not a subgroup). We can now define the basic module that will repeat through the hierarchy. An HW-module consists of three elements: (*t, G*_0_, *P*). The output of an HW-module is *μ*(*I*) = *P*_*gϵG*_0 (*(I, gt)*). Note that if *G*_0_ is a set of translations and *P*_*g*_(*·*) = max_*g*_(*·*), then one such HW-module is exactly equivalent to an HMAX C-unit (defined in the main text). The subset *G*_0_ can be thought of as the pooling domain. In the case of translation it has the same interpretation as a spatial region as in HMAX.

Consider, for simplicity, the case of 1D images (centered in zero) transforming under the 1D locally compact group of translations. What are the conditions under which an HW-module will be invariant over the range *G*_0_ = [*b, b*]? Let *P*_*g*_(·) := ∑_*xϵ[−b, b]*_ *η(·)*, where *η* is a positive, bijective function. The signature vector components will be given by

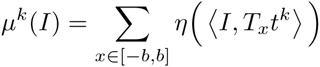

where *T*_*x*_ is the operator acting on a function *f* as *T*_*x*_*f* (*x*^*t*^) = *f* (*x*^*t*^ - *x*). Suppose we transform the image *I* (or equivalently, the template) by a translation of 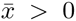, implemented by 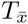. Under what conditions does *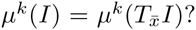* Note first that *I, T*_*x*_*t*^*k*^ = (*I * t*^*k*^)(*x*), where *\** indicates convolution. By the properties of the convolution operator, we have 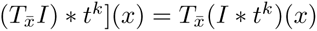 which implies

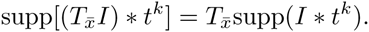

This observation allows us to write a condition for the invariance of the signature vector components with respect to the translation *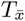* (see also Fig. 7). For a positive nonlinearity *η*, (no cancelations in the sum) and bijective (the support of the dot product is unchanged by applying *η*) the condition for invariance is:

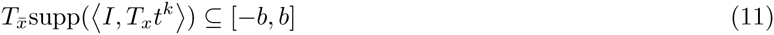

**Figure 7:**
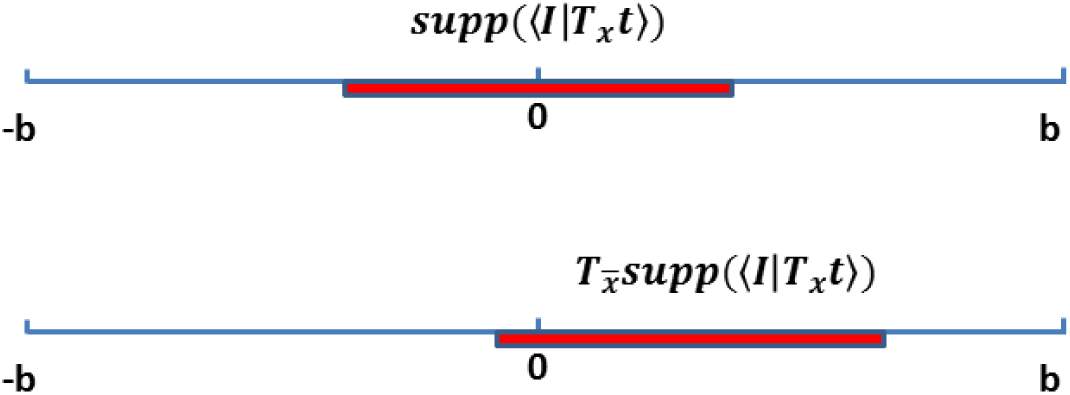
Localization condition of the S-unit response for invariance under the transformation *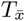*

Eq. 11 is a localization condition on the S-unit response. It is necessary and sufficient for invariance. In this case, eq. (9) is trivial since we are considering group transformations.

#### 5.2 The second regime: class-specific invariance

So far, we have explained how the localization properties of the S-response allow invariance in the case of partially observed group transformations. Next, we show how localization still enables approximate invariance (*E*-invariance) even in the case of non-group (smooth) transformations. However, as will be shown below, in order for eq. (9) to be (approximately) satisfied, the class of templates needs to be much more constrained than in the group case.

Consider a smooth transformation parametrized by *r ϵ* ℝ, *T*_*r*_; the Taylor expansion of *T*_*r*_*I* w.r.t. *r* around, e.g., zero is:

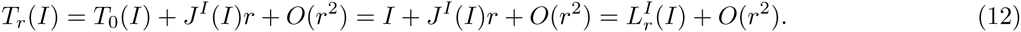

where *J*^*I*^ is the Jacobian of the transformation *T*, and *L*^*I*^ (*·*) = *e*(*·*) + *J* ^*I*^ (*·*)*r*. The operator *L*^*I*^ corresponds to the best linearization around the point *r* = 0 of the transformation *T*_*r*_. Let *R* be the range of the parameter *r* such that *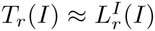.* If the localization condition holds for a subset of the transformation parameters contained in *R*, i.e.

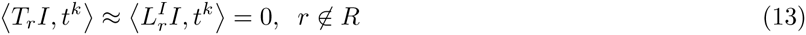

and as long as the pooling range *P*, in the *r* parameter is chosen so that *P ⊆ R*, then we are back in the group case, and the same reasoning used above for translation still holds.

However this is not the case for eq. (9). The tangent space of the image orbit is given by the Jacobian, and it clearly depends on the image itself. Since the tangent space of the image and of the template will generally be different (see Fig. 8), this prevents eq. (9) from being satisfied. More formally, for *r ϵ R*:

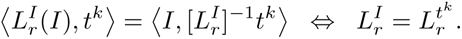

That is, eq. (9) is only satisfied when the image and template “transform the same way” (see Fig. 8).

**Figure 8:**
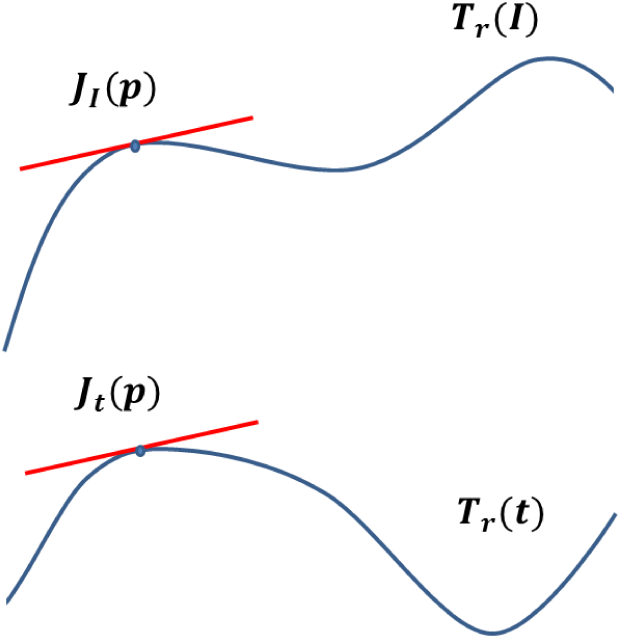
The Jacobians of the orbits of the image around the point *p* and the template must be approximately equal for eq. (9) to hold in the case of smooth transformations.

To summarize, the following three conditions are needed to have invariance for non-group transformations:

1. The transformation must be differentiable (the Jacobian must exist).
2. A localization condition of the form in eq. (13) must hold to allow a linearization of the transformation.
3. The image and templates must transform ”in the same way”, i.e. the tangent space of their orbits (in the localization range) must be equal. This is equivalent to *J* ^*I*^ ≡ *J* ^*tk*^.

**Remark**: The exposition of the theory given here is specialized for the relevant case of the general theory. In general, we allow each “element” of the signature (as defined here) to be a vector representing a distribution of one-dimensional projections of the orbit. See [37] for details.

### 6 Illumination invariance

**Figure 9:**
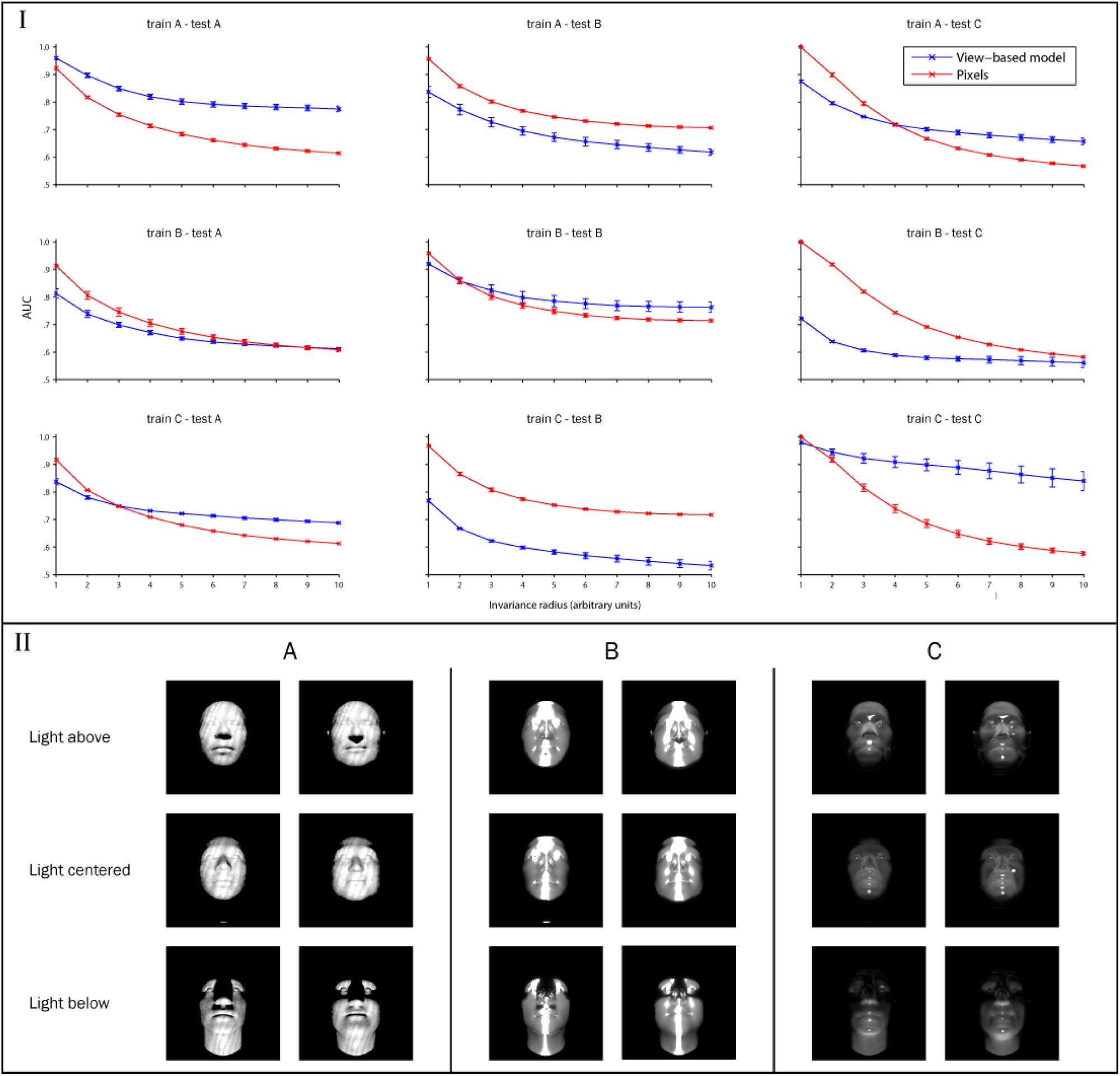
Class-specific transfer of illumination invariance. Bottom panel (II): Example images from the three classes. Top panel (I): The left column shows the results of a test of illumination invariance on statues of heads made from different materials (class A), the middle column shows results for class B and the right column shows the results for class C. The view-based model (blue curve) was built using images from class A in the top row, class B in the middle row, and class C in the bottom row. The abscissa of each plot shows the maximum invariance range (arbitrary units of the light source’s vertical distance from its central position) over which target and distractor images were generated. The view-based model was never tested on any of the images that were used as templates. Error bars (+/- one standard deviation) were computed over 20 cross validation runs using different choices of template and test images.

Illumination is also a class-specific transformation. The appearance of an object after a change in lighting direction depends both on the object’s 3D structure and on its material properties (e.g. reflectance, opacity, specularities). Figure 9 displays the results from a test of illumination-invariant recognition on three different object classes which can be thought of as statues of heads made from different materials—A: wood, B: silver, and C: glass. The results of this illumination-invariance test follow the same pattern as the 3D rotation-invariance test. In both cases the view-based model improves the pixel-based models’ performance when the template and test images are from the same class (fig. 9—plots on the diagonal). Using templates of a different class than the test class actually lowered performance below the pixel-based model in some of the tests e.g. train A–test B and train B–test C (fig. 9—off diagonal plots). This simulation suggests that these object classes have high 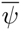 respect to illumination transformations. However, the weak performance of the view-based model on the silver objects indicates that it is not as high as the others (see the table below). This is because the small differences in 3D structure that define individual heads give rise to more extreme changes in specular highlights under the the transformation.

**Table 2:**
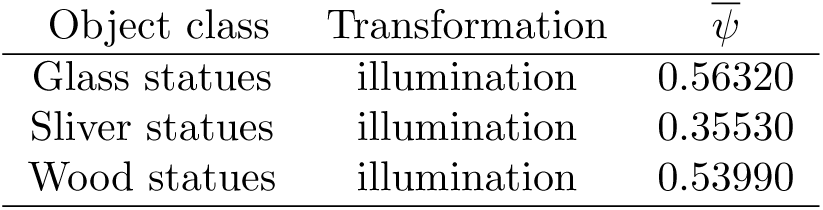
Table of illumination transformation compatibilities

| Object class | Transformation | $\bar{\psi}$ |
| --- | --- | --- |
| Glass statues | illumination | 0.56320 |
| Sliver statues | illumination | 0.35530 |
| Wood statues | illumination | 0.53990 |

### 7 Pose-invariant body recognition

**Figure 10:**
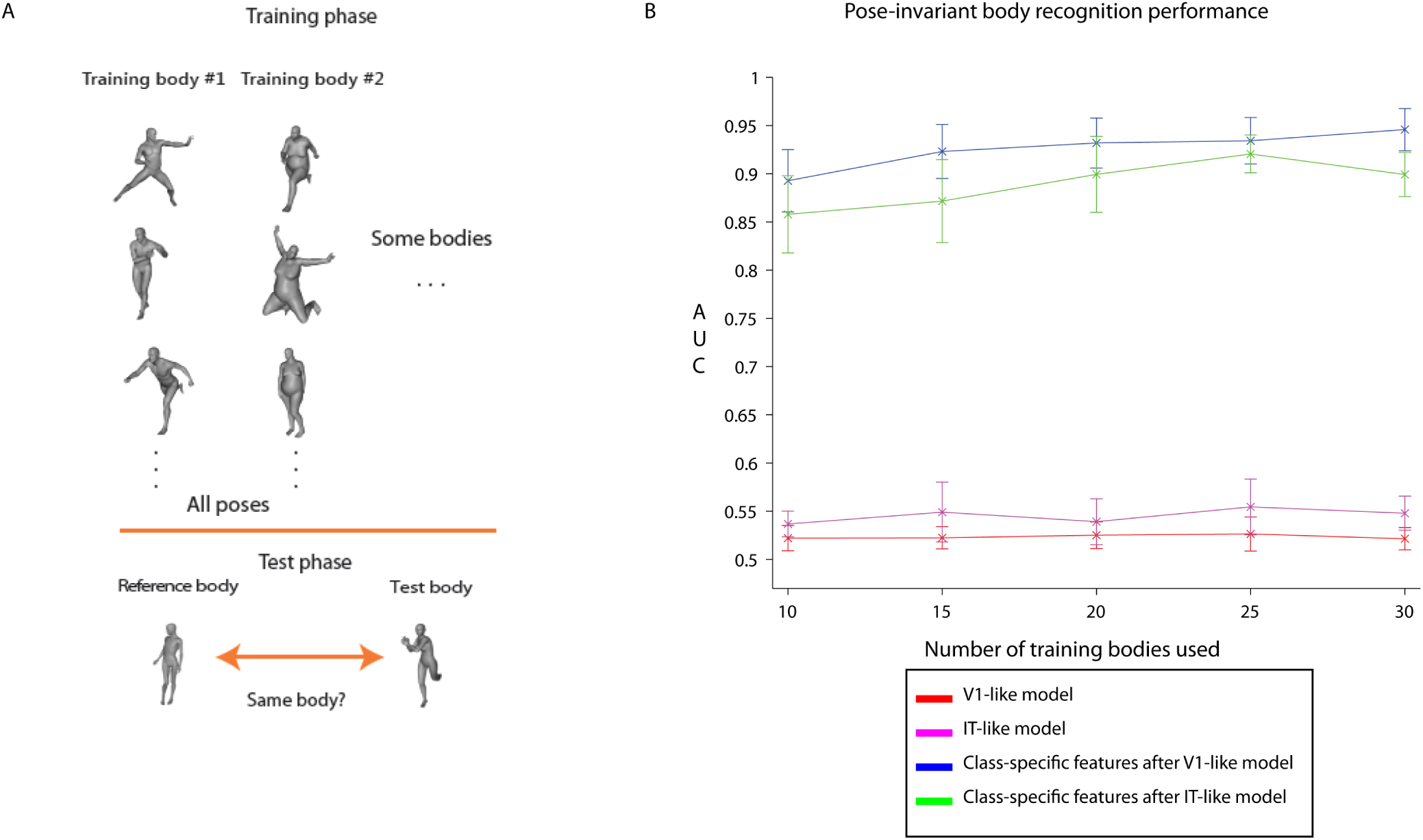
**A.** Example images for the pose-invariant body-recognition task. The images appearing in the training phase were used as templates. The test measures the model’s performance on a same-different task in which a reference image is compared to a query image. ‘Same’ responses are marked correct when the reference and query image depict the same body (invariantly to pose-variation). **B.** Model performance: area under the ROC curve (AUC) for the same-different task with 10 testing images. The X-axis indicates the number of bodies used to train the model. Performance was averaged over 10 cross-validation splits. The error bars indicate one standard deviation over splits.

Let *B* = *{b*_1_, *b*_2_, *…, b*_*n*_} be a set of bodies and *P* = *{p*_1_, *p*_2_, *…, p*_*n*_} be a set of poses. Let *d* be the dimensionality of the images. We define the rendering function *t*_*p*_ : *B* ⟶ ℝ*d.* In words, we say *t*_*p*_[*b*] renders an image of body *b* in pose *p*. In that case the argument *b* is the template and the subscript *p* indicates the transformation to be applied.

We obtain the signature vector *μ* : *X* ⟶ ℝ^*m*^ by pooling the inner products of the input image with different renderings of the same template.

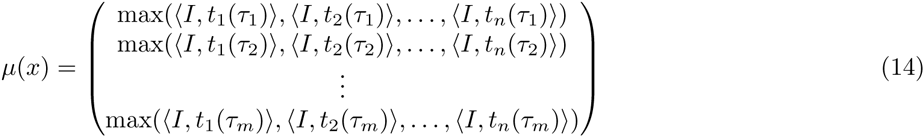

As in some HMAX implementations (e.g., Serre et al. (2007) [120]), we used a Gaussian radial basis function for the S-unit response. It has similar properties to the normalized dot product.

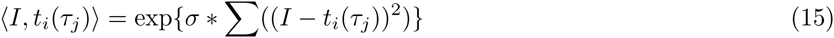

Where *σ* is the Gaussian’s variance parameter.

The class-specific layer takes in any vector representation of an image as input. We investigated two hierarchical architectures built off of different layers of the HMAX model (C1 and C2-global) [120]—referred to in fig. 10 as the V1-like and IT-like models respectively.

For the pose-invariant body recognition task, the template images were drawn from a subset of the 44 bodies—rendered in all poses. In each of 10 cross-validation splits, the testing set contained images of 10 bodies that never appeared in the model-building phase—again, rendered in all poses (fig. 10).

The HMAX models perform almost at chance. The addition of the class-specific mechanism significantly improves performance on this difficult task. That is, models without class-specific features were unable to perform the task while class-specific features enabled good performance on this difficult invariant recognition task (fig. 10).

Downing and Peelen (2011) argued that the extrastriate body area (EBA) and fusiform body area (FBA) “jointly create a detailed but cognitively unelaborated visual representation of the appearance of the human body”. These are perceptual regions—they represent body shape and posture but do not explicitly represent high-level information about “identities, actions, or emotional states” (as had been claimed by others in the literature [121]). The model of body-specific processing suggested by the simulations presented here is broadly in agreement with this view of EBA and FBA’s function. It computes, from an image, a body-specific representation that could underlie many further computations e.g. action recognition, emotion recognition, etc.

### 8 Supplementary simulations on the development of domain-specific regions

**Figure 11:**
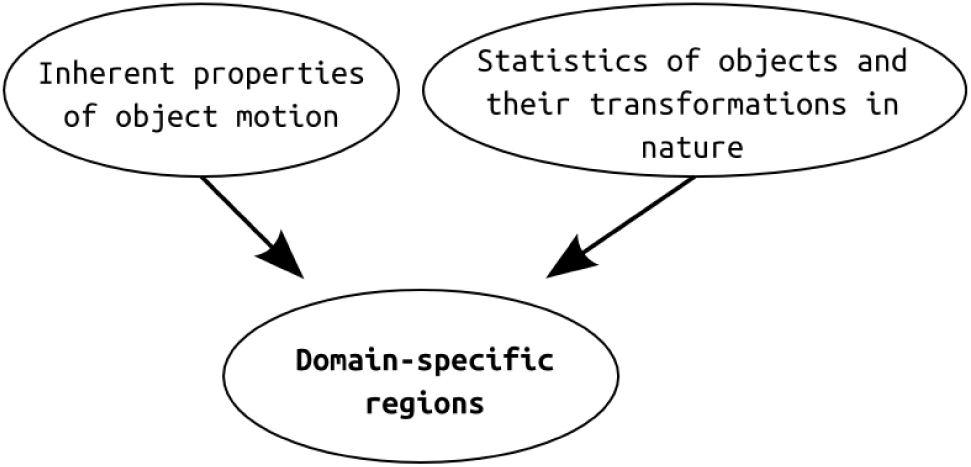
Two factors are conjectured to influence the development of domain-specific regions.

We consider three different arbitrary choices for the distributions of objects from five different categories: faces, bodies, vehicles, chairs, and animals (see table 3). Importantly, one set of simulations used statistics which were strongly biased against the appearance of faces as opposed to other objects.

**Table 3:** Numbers of objects used for each simulation. In the “realistic” simulation, there were proportionally more faces.

|  | Name of simulation | Faces | Bodies | Animals | Chairs | Vehicles |
| --- | --- | --- | --- | --- | --- | --- |
| A. | “Realistic” | 76 | 32 | 16 | 16 | 16 |
| B. | Uniform | 30 | 30 | 30 | 30 | 30 |
| C. | Biased against faces | 16 | 32 | 36 | 36 | 36 |

**Figure 12:**
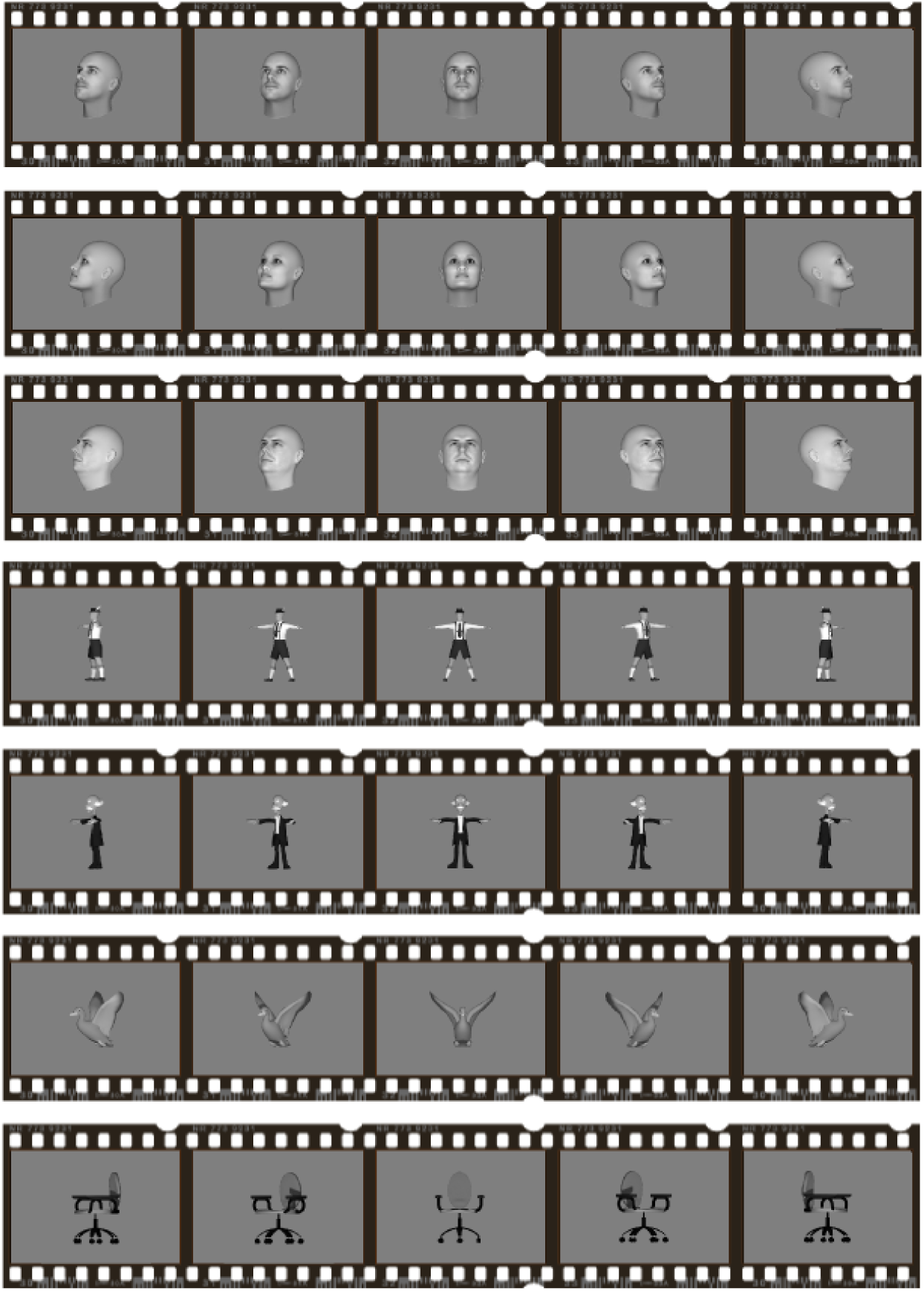
Example object videos (transformation sequences) used in the *Ψ*-based clustering experiments.

**Figure 13:**
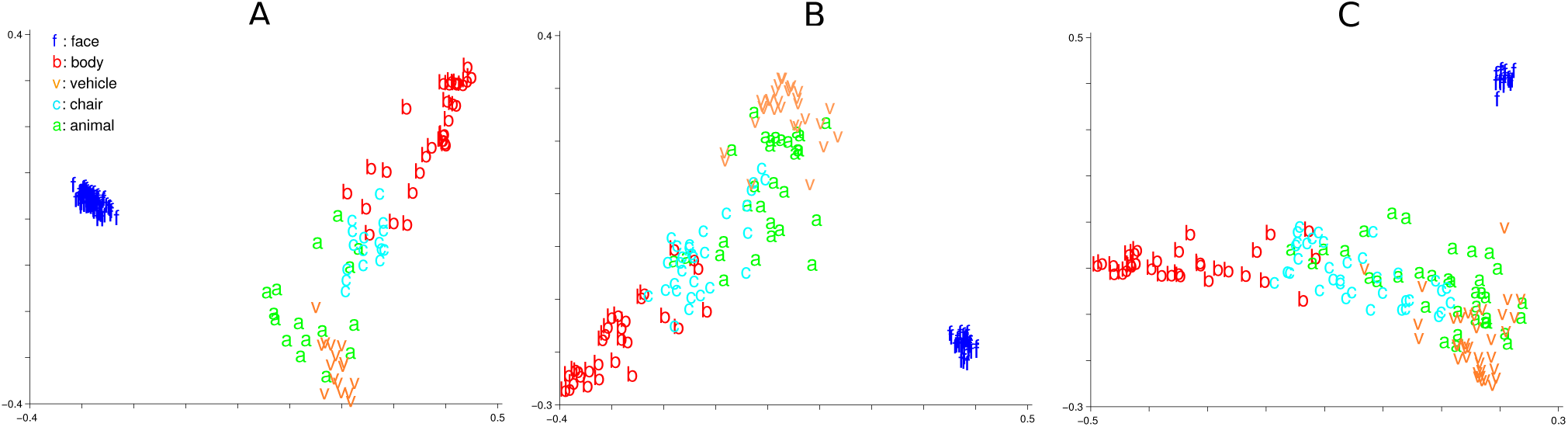
Multidimensional Scaling (MDS) [114] visualizations of the object sets under the *Ψ*(*A, B*)-dissimilarity metric for the three object distributions: A. “realistic”, B. uniform, and C. biased against faces (see table 3).

**Figure 14:**
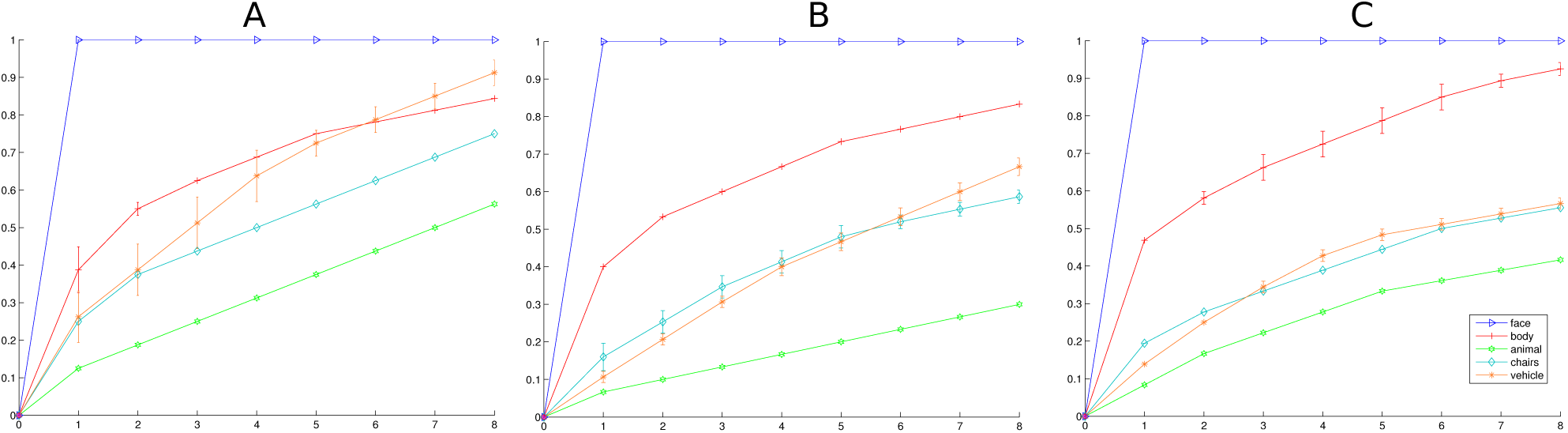
The percentage of objects in the first N clusters containing the dominant category object (clusters sorted by number of objects in dominant category). A, B and C are respectively, the “realistic” distribution, uniform distribution, and the biased against faces distribution (see table 3)). 100% of the faces go to the first face cluster—only a single face cluster developed in each experiment. Bodies were more “concentrated” in a small number of clusters, while the other objects were all scattered in many clusters—thus their curves rise slowly. These results were averaged over 5 repetitions of each clustering simulation using different randomly chosen objects.

**Figure 15:**
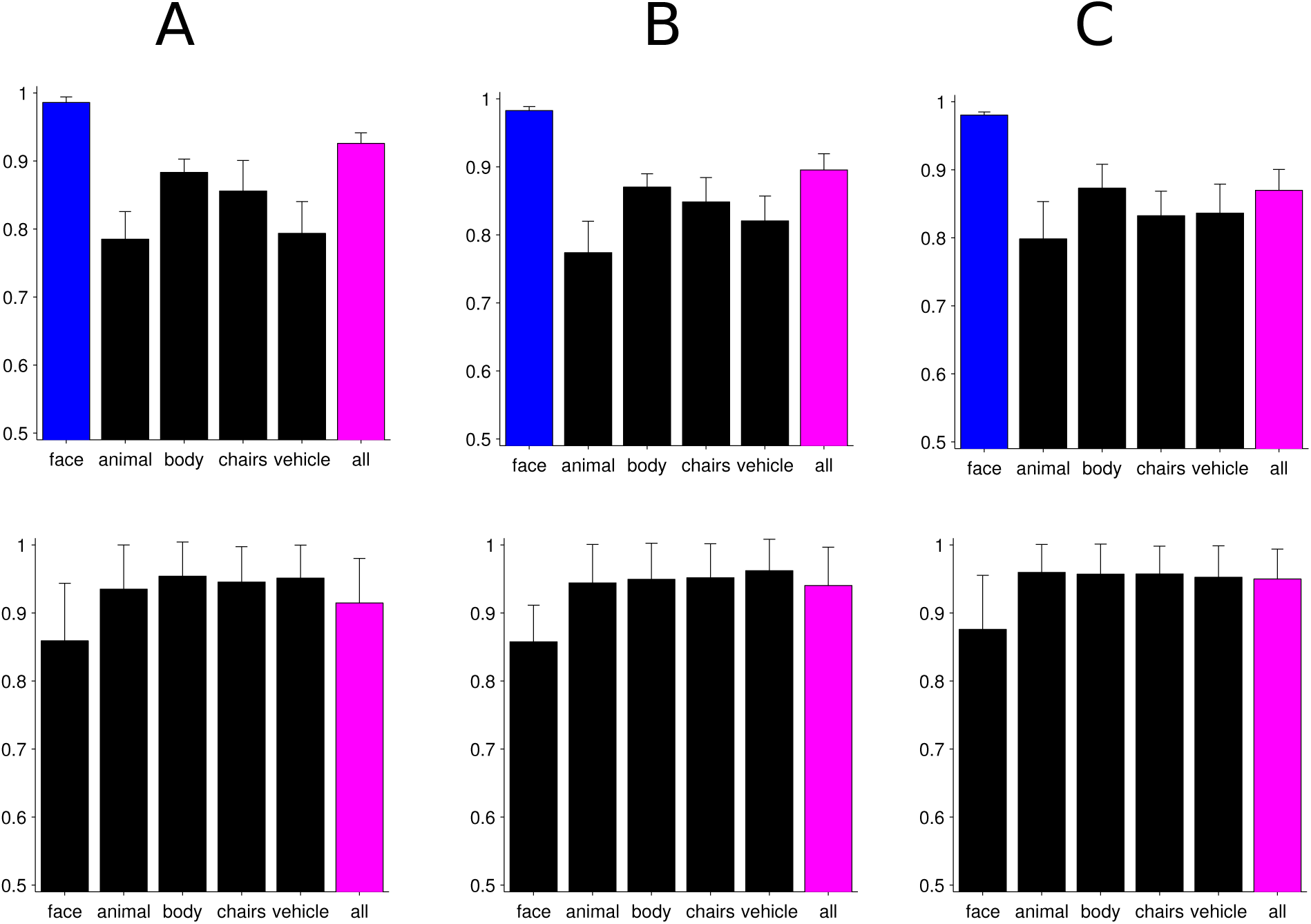
The classification performance on face recognition, a subordinate-level task (top row) and car vs. airplane, a basic-level categorization task (bottom row) using templates from each cluster. 5-fold cross-validation, for each fold, the result from the best-performing cluster of each category is reported. A, B and C indicate “realistic”, uniform, and biased distributions respectively (see table 3). Note that performance on the face recognition task is strongest when using the face cluster while the performance on the basic-level car vs. airplane task is not stronger with the vehicle cluster (mostly cars and airplanes) than the others.

### 9 Supplementary methods

#### Stimuli

##### Illumination

Illumination: Within each class the texture and material properties were exactly the same for all objects. We used Blender to render images of each object with the scene’s sole light source placed in different locations. The 0 position was set to be in front of the object’s midpoint; the light was translated vertically. The most extreme translations brought the light source slightly above or below the object. Material data files were obtained from the Blender Open Material Repository (http://matrep.parastudios.de/). 40 heads were rendered with each material type. For each repetition of the experiment, 20 were randomly chosen to be templates and 20 to be testing objects. Each experiment was repeated 20 times with different template and testing sets.

##### Bodies / pose

DAZ 3D Studio was used to render each of 44 different human bodies under 32 different poses, i.e., 44*32 =1408 images in total.

#### Body-pose experiments

For the body-pose invariance experiments (fig. 10), the task was identical to the test for unfamiliar faces and novel object classes. The same classifier (Pearson correlation) was used for this experiment. Unlike rotation-in-depth, the body-pose transformation was not parameterized.

#### Clustering by transformation compatibility

Pseudocode for the clustering algorithm is given below (algorithm 1).

Let *A*_*i*_ be the *i*_*th*_ frame of the video of object A transforming and *B*_*i*_ be the *i*_*th*_ frame of the video of object B transforming. The Jacobian can be approximated by the “video” of difference images: *J*_*A*_(*i*) = *|A*_*i*_ - *A*_*i*+1_*|* (*∀i*). The “instantaneous” transformation compatibility is *Ψ*(*A, B*)(*i*) := *(J*_*A*_(*i*), *J*_*B*_(*i*)). Thus for a range of parameters *i ϵ R* = [*-r, r*], the empirical transformation compatibility between *A* and *B* is

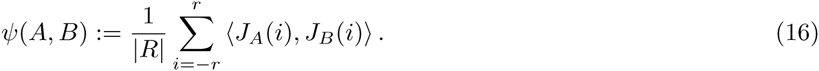

The transformation compatibility 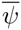 of a cluster *C* was defined as the average of the pairwise compatibilities *Ψ*(*A, B*) of all objects in *C*.

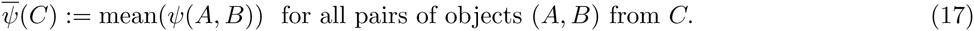

##### Algorithm 1 Iterative clustering algorithm interpreted as a model of ventral stream development

**Input:** All Objects: *O*, *i*_*th*_ Object: *O*_*i*_ where *i* = 1*…N*, Threshold: *T*)

~~~
**Output:** ClusterLabels
ClusterLabels(1) = 1
*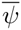* = computeCompatibility(ClusterLabels)
**for** *i* = 2 **to** *N* **do**
    Ψ = computeCompatibilityWithEveryCluster(*i, O*, ClusterLabels)
    [MaxValue MaxIndex] = max(*Ψ*)
    **if** MaxValue *> T* **then**
       ClusterLabels(*i*) = MaxIndex //Assign to the cluster with the highest compatibility.
    **else**
       ClusterLabels(*i*) = max(ClusterLabels) + 1 //Create a new cluster
    **end if**
    *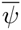* = updateCompatibility(*Ψ*, CurrentClusterCompatibility, ClusterLabels(*i*))
**end for**
**Function**   computeCompatibilityWithEveryCluster(IDX,AllObjects,ClusterLabels)
//Initialize *Ψ* as an empty array of length #Clusters.
**for** *i* = 1 **to** #Clusters **do**
    Objects = GetObjectsFromCluster(*i*, AllObjects, ClusterLabels)
    **for** *j* = 1 **to** #Objects **do**
       tmpArray(*j*) = compatibilityFunction(AllObjects(IDX), Objects(*j*))
    **end for**
    Ψ(*i*) = mean(tmpArray);
**end for**
Return *Ψ*
**EndFunction**
~~~

## Acknowledgments

We would like to thank users bohemix and SoylentGreen of the Blender Open Material Repository for contributing the materials used to create the images for the illumination simulations (in supplementary information). We also thank Andrei Rusu, Leyla Isik, Chris Summerfield, Winrich Freiwald, Pawan Sinha, and Nancy Kanwisher for their comments on early versions of this manuscript, and Heejung Kim for her help preparing one of the supplementary figures. This report describes research done at the Center for Biological & Computational Learning, which is in the McGovern Institute for Brain Research at MIT, as well as in the Dept. of Brain & Cognitive Sciences, and which is affiliated with the Computer Sciences & Artificial Intelligence Laboratory (CSAIL). This material is based upon work supported by the Center for Brains, Minds, and Machines (CBMM), funded by NSF STC award CCF-1231216. This research was also sponsored by grants from the National Science Foundation (NSF-0640097, NSF-0827427), and AFOSR-THRL (FA8650-05-C-7262). Additional support was provided by the Eugene McDermott Foundation.

